# Genomic characterization and curation of UCEs improves species tree reconstruction

**DOI:** 10.1101/828335

**Authors:** Matthew H. Van Dam, James B. Henderson, Lauren Esposito, Michelle Trautwein

**Affiliations:** Entomology Department, Institute for Biodiversity Science and Sustainability, California Academy of Sciences, 55 Music Concourse Dr., San Francisco, CA, 94118 USA; Center for Comparative Genomics, Institute for Biodiversity Science and Sustainability, California Academy of Sciences, 55 Music Concourse Dr., San Francisco, CA, 94118 USA

**Author notes:** Correspondence to be sent to: Matthew H. Van Dam, Entomology Department, Institute for Biodiversity Science and Sustainability, California Academy of Sciences, 55 Music Concourse Dr., San Francisco, CA, 94118 USA.

**Keywords:** Ultraconserved elements, species tree, gene tree, phylogenomic markers, bait capture, ASTRAL, conserved nonexonic elements, exon capture, anchored hybrid enrichment, Hymenoptera, Coleoptera, carangimorph, ants, weevils, mammal, songbird

## Abstract

Ultraconserved genomic elements (UCEs), are generally treated as independent loci in phylogenetic analyses. The identification pipeline for UCE probes is agnostic to genetic identity, only selecting loci that are highly conserved, single copy, without repeats, and of a particular length. Here we characterized UCEs from 12 phylogenomic studies across the animal tree of life, from birds to marine invertebrates. We found that within vertebrate lineages, UCEs are mostly intronic and intergenic, while in invertebrates, the majority are in exons. We then curated 4 different sets of UCE markers by genomic category from 5 different studies including; birds, mammals, fish, Hymenoptera (ants, wasps and bees) and Coleoptera (beetles). Of genes captured by UCEs, we find that many are represented by 2 or more UCEs, corresponding to non-overlapping segments of a single gene. We considered these UCEs to be non-independent, merged all UCEs that belonged to a particular gene, constructed gene and species trees, and then evaluated the subsequent effect of merging co-genic UCEs on gene and species tree reconstruction. Average bootstrap support for merged UCE gene trees were significantly improved across all datasets. Increased loci length appears to drive this increase in bootstrap support. Additionally, we found that gene trees generated from merged UCEs were more accurate than those generated by unmerged and randomly merged UCEs, based on our simulation study. This modest degree of UCE characterization and curation impacts downstream analyses and demonstrates the advantages of incorporating basic genomic characterizations into phylogenomic analyses.

## INTRODUCTION

Phylogenomic methods rely on sampling orthologous loci from the genomes of non-model organisms and then using these loci for the estimation of evolutionary relationships. Commonly used sampling strategies include; ultraconserved genomic elements (UCEs), *sensu* Faircloth et al. 2012, anchored hybrid enrichment (Lemmon et al. 2012), exon capture (Bi et al. 2013), transcriptomes, along with homologous *k*-mer blocks (Sanderson et al. 2017), and conserved nonexonic elements (CNEEs; Edwards et al. 2017). While UCEs and anchored hybrid enrichment markers are generally identified without regard to what genomic class they fall into, transcriptomes, exon capture (Bi et al. 2013) and CNEEs (Edwards et al. 2017) each select for a specific class of marker as their names entail.

UCEs are among the most widely used types of phylogenomic markers, and have been used to resolve both higher level and population level phylogenetic relationships (McCormack et al. 2012, Winker et al. 2018). They are found throughout the tree of life, including Cnidaria, flat worms (Platyhelminthes), arachnids, insects, as well as in birds and mammals (Quattrini et al. 2018, Locke et al. 2018, Faircloth 2017, Van Dam et al. 2018, Moyle et al. 2016, Esselstyn et al. 2017). UCEs are beneficial for their ease of capture from non-model organisms. In addition, UCEs and other probe based approaches provide access to loci from degraded DNA in museum specimens (Bi et al. 2013, McCormack et al. 2016, Blaimer et al. 2016, Van Dam et al. 2017). The process of identifying UCEs is agnostic to the genetic identity of each UCE and instead simply selects for loci that are highly conserved, single copy, lack repeats (and having a user defined GC content), and are of a particular length (typically 160bp length) (Faircloth et al. 2012; Faircloth 2017).

Independent of phylogenetics, a large volume of research has been conducted on identifying the function of UCEs in the genome and towards understanding why they are highly conserved over many millions of years (Dermitzakis et al. 2003; Bejerano et al. 2004; Sandelin et al. 2004; Woolfe et al. 2004; Vavouri et al. 2007; McCole et al. 2014, 2018; Kushawah and Mishra 2017). The sets of UCEs used in phylogenomics have not been formally documented as to their genomic categorization even at the basic level (intronic, exonic and or intergenic) (but see Jarvis et al. 2014). The genomic identity of phylogenetic markers affects how data is treated in analyses (e.g. models of nucleotide substitution rate based on codon position), as well as their potential phylogenetic informativeness (Gilbert el al. 2018). Generally, UCEs are considered primarily non-coding entities and are treated as such in phylogenetic analyses (but see Jarvis et al. 2014, Bossert et al. 2018, Branstetter et al, 2017b).

Like many different classes of phylogenomic markers, UCEs have long been treated as anonymous, independent loci. A markers’ independence is relevant to subsequent analyses– particularly in multi-species coalescent tree estimation. Treating non-independent samples as independent violates the assumption of statistical independence between samples which could bias results by giving more representation to particular gene tree topologies (see Szöllősi et al. 2015).

Here we examined 12 sets of UCEs that have been used for phylogenomic estimation across the tree of life and identify the genetic class of each UCE as intronic, exonic, or intergenic. We examined how these characterizations compare between organismal classes. We considered 5 previous UCE studies in mammals, birds, fish, and insects (Hymenoptera: wasps, ants and bees, and Coleoptera: beetles) more extensively and examined the phylogenetic utility of intergenic and genic UCEs across taxa. Our data exploration revealed that many genes are actually represented by multiple non-overlapping UCEs (referred to throughout as co-genic UCEs in accordance with Scornavacca and Galtier, 2017). Co-genic UCEs that are then used to generate multiple gene trees from the same gene violate the assumption of independence between loci in subsequent multi-species coalescent analyses. We then concatenated (referred to herein as merged) all co-genic UCEs and performed phylogenetic reconstructions of gene and species trees to examine the effect of merging co-genic UCEs on average bootstrap support values (ABS). Finally, we performed a simulation study to test the effects of merging co-genic UCEs on the accuracy of phylogenetic inference of gene trees and species trees.

## METHODS

### Genomic Characterization of UCEs across the Tree of Life

First, we characterized the genomic identity of 12 sets of UCEs representing diverse regions of the tree of life from mammals to marine invertebrates (Figs. 1–2, Table 1). We made our characterizations by blasting (using *blastn* version 2.9.0, Camacho et al. 2008) the base taxon’s probes of each taxon set back to their base genome (the genome to which all other genomes were aligned to in the UCE probe set design). For the tetrapod and acanthomorph fish UCE sets, a base genome was not used in probe design and instead probes were identified by performing an all to all alignment (Faircloth et al. 2012). For these sets, we assigned a new base genome as the taxon for which the most UCE loci were recovered: chicken (*Gallus gallus)* for tetrapods and medaka (*Oryzias latipes)* for the acanthomorph fish probe sets (Faircloth et al. 2012; Faircloth et al. 2013). For all 12 UCE sets, we downloaded each original probe set, identified and extracted the base-genome’s (or assigned base genome’s) specific probes from the total probe fasta file and generated a new fasta containing just the base-genome’s probes. Then we blasted these base genome probes against the base-genome using *blastn* (Camacho et al. 2008). The resulting m8 file from the *blastn* search was then filtered for 100% matches over the 120bp length of the probe.

**Figure 1.**
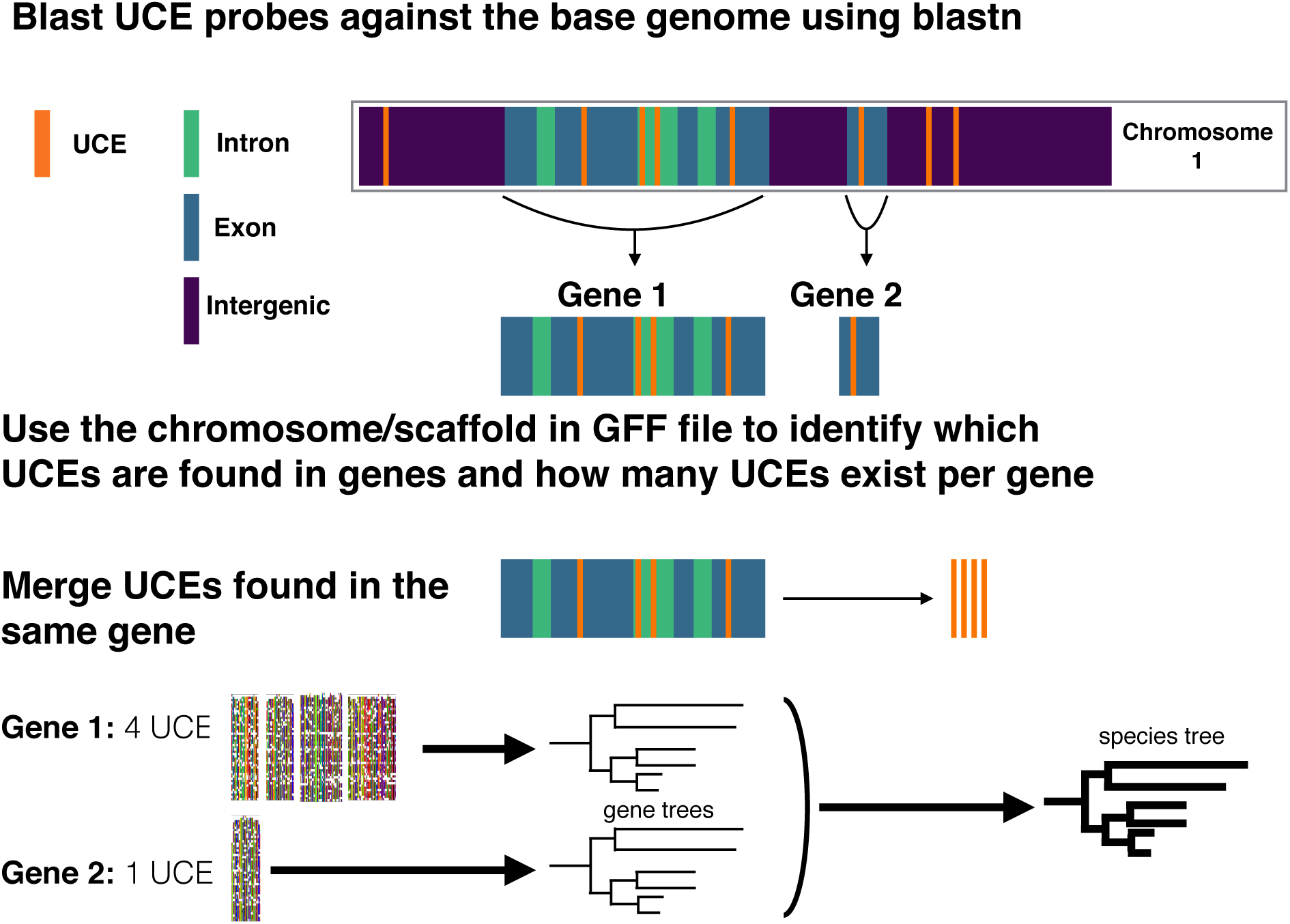
General workflow used to identify co-genic UCEs and merge them for gene tree and species tree reconstruction.

**Figure 2.**
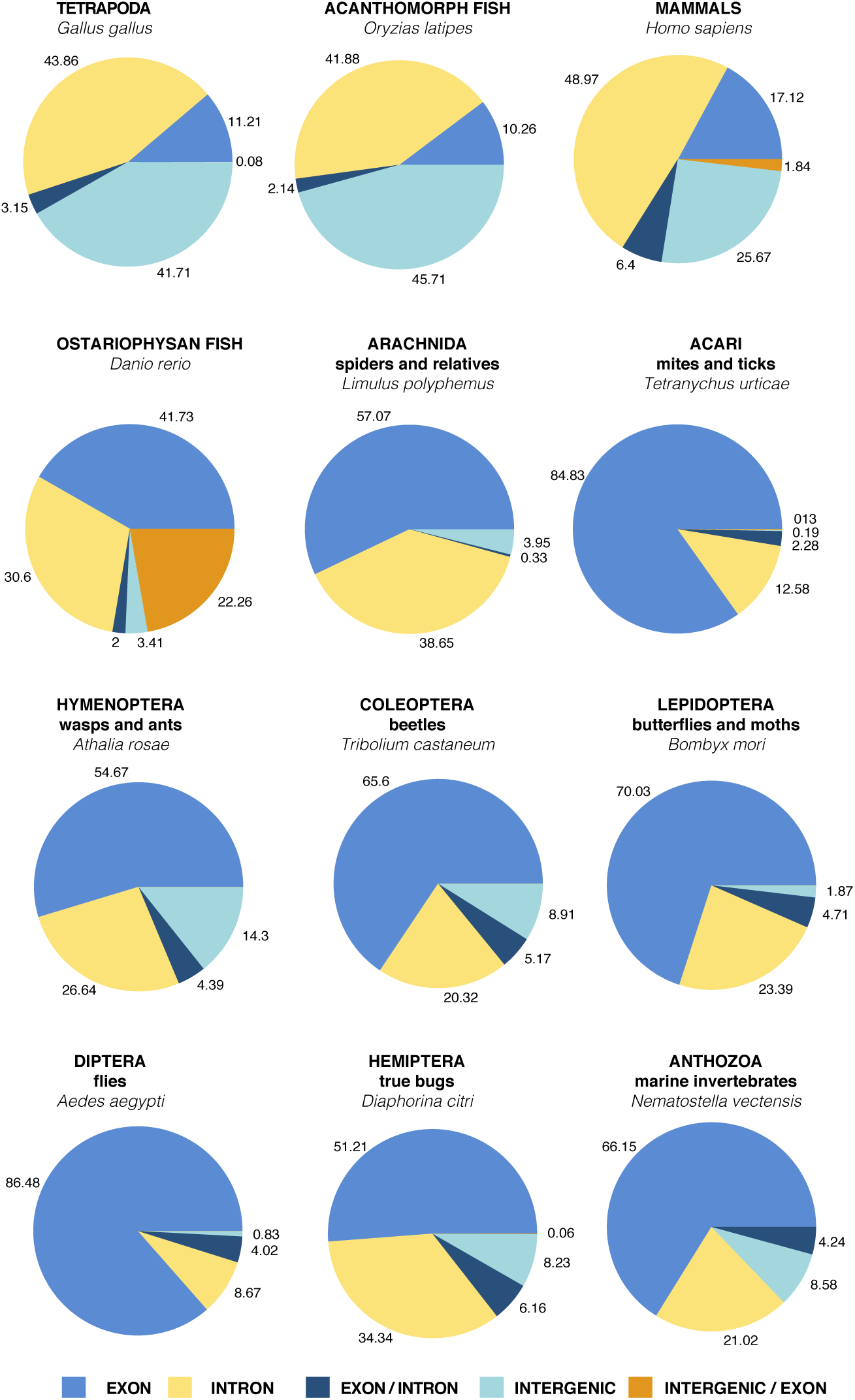
Characterization of 12 UCE probe sets according to the annotated base genome for each set. UCEs are put into 5 different categories: intergenic (not in a gene), exon, intron, exon/intron, and intergenic/exon. The categories exon/intron and intergenic/exon represents those UCEs that span an intron and an exon or an exon and an intergenic region.

**TABLE 1.** Characterization of 12 UCE probe sets according to the annotated base genome (in the base taxon column) for each set.

| UCE set | BASE TAXON | EXON | INTRON | EXON/ INTRON | INTER- GENIC | INTER- GENIC/ EXON | TOTAL | CITATION |
| --- | --- | --- | --- | --- | --- | --- | --- | --- |
| Tetrapod-5K | *Gallus gallus* | 563 | 2203 | 158 | 2095 | 4 | 5023 | Faircloth et al. 2012 |
| Acantho- morphs-1K | *Oryzias latipes* | 134 | 547 | 28 | 597 | 0 | 1306 | Faircloth et. al. 2013 |
| Ostariophysa ns-2.7K | *Danio rerio* | 1125 | 825 | 54 | 92 | 600 | 2696 | Faircloth et al. 2018 |
| Anthozoan_V 1-UCE-only | *Nematostella vectensis* | 686 | 218 | 0 | 89 | 44 | 1037 | Quattrini et al. 2017 |
| Arachnida- 1.1K | *Limulus polyphemus* | 347 | 235 | 2 | 24 | 0 | 608 | Faircloth 2017 |
| Acari-1.8K | *Tetranychus urticae* | 1342 | 199 | 36 | 3 | 2 | 1582 | Van Dam et al. 2018 |
| Hymenoptera- v2 | *Athalia rosae* | 1170 | 570 | 94 | 306 | 0 | 2140 | Branstetter et al. 2017b |
| Coleoptera- 1.1K | *Tribolium castaneum* | 736 | 228 | 58 | 100 | 0 | 1122 | Faircloth 2017 |
| Lepidoptera- 1.3K | *Bombyx mori* | 937 | 313 | 63 | 25 | 0 | 1338 | Faircloth 2017 |
| Diptera-2.7K | *Aedes aegypti* | 1766 | 177 | 82 | 17 | 0 | 2042 | Faircloth 2017 |
| Hemiptera- 2.7K | *Diaphorina citri* | 865 | 580 | 104 | 139 | 1 | 1689 | Faircloth 2017 |
| Mammals | *Homo sapiens* | 3450 | 9871 | 1290 | 5175 | 371 | 20157 | Van Dam et al. in prep |

To identify a UCE’s position within a genome, we identified the UCE’s scaffold and/or chromosome and position from the m8 file. The scaffold and/or chromosome and position information allowed us to search the base-genome’s GFF file to identify overlap between each UCE and particular gene features. The positions of UCEs that fell within introns were inferred from start and stop positions of exons within gene regions. The NCBI records for the specific genomes and GFF files are listed in Supplementary Material S1.

### Focal taxa UCE characterization and curation

#### Data acquisition and alignment

To examine how UCE characterization can affect phylogenetic inference, we more extensively considered UCE data from 5 different phylogenetic studies using 4 different UCE bait sets representing: **weevils** (Coleoptera UCE baits: Faircloth 2017; Van Dam et al. 2017), **ants** (Hymenoptera UCE baits-V2:Branstetter et al. 2017b), **mammals** (Tetrapod 5K-UCE baits: Faircloth et al. 2012; Esselstyn et al. 2017), **songbirds** (Tetrapod 5K-UCE baits: Faircloth et al. 2012; Moyle et al. 2016) and carangimorph **fish** (acanthomorph fish UCE baits: Faircloth et al. 2013; Harrington et al. 2016), see Table 2. Using the original data from these studies, we followed their assembly and matrix construction procedures largely using the *PHYLUCE* pipeline (Faircloth et al. 2012; Faircloth 2016).

**Table 2.** Taxa column lists the previous studies used here. Second column lists the number of individual taxa used in each study, followed by the total number of UCEs. Last column lists the minimum number of taxa allowed in a UCE locus for it to be included/average number of taxa per UCE locus.

| TAXA | # of TAXA | Total # of UCEs | Minimum # of TAXA Per Locus /Average # of TAXA |
| --- | --- | --- | --- |
| **weevils** (Van Dam et al. 2017) | 48 | 368 | 33/38 |
| **ants** (Branstetter et al. 2017b) | 101 | 2571 | 10/89 |
| **birds** (Moyle et al. 2016) | 106 | 4347 | 79/102 |
| **mammals** (Esselstyn et al. 2017) | 100 | 3594 | 70/88 |
| **carangimorph fish** (Harrington et al. 2016) | 55 | 1014 | 41/52 |

For three of the datasets (fish, ants and weevils), aligned fasta sequences were already available. For birds and mammals, aligned reads were not available, so we downloaded raw reads and followed the procedures taken by the previous authors to create our aligned matrices. In the case of the mammal dataset (Esselstyn et al. 2017), we downloaded the raw reads and also extracted UCE loci from the same genomes used by the authors, and then combined these data.

For all alignments, we used the *R* package *ips* (Heibl 2008) and removed any ragged ends with the function “*trimEnds*” having a minimum of 4 taxa present in the alignment and filled any gap character “-” with “n” before the first and last non-ambiguous nucleotide.

### Curation of genic UCEs

After determining which UCEs were found within genes from our focal UCE sets, we curated sets of genic and intergenic UCEs for weevils, ants, songbirds, mammals and caragimorph fish, (from here on referred to as **Genic** and **Intergenic)**. Next, across our taxa sets we identified genes that contain multiple UCEs (co-genic UCEs) (Table 3). Co-genic UCEs were then merged into a single alignment per gene using scripts we developed (Supplementary Material). We then curated 2 sets of UCEs for each taxon group: one that included all UCEs-called **Unmerged** and another that included all merged co-genic UCEs + all remaining UCEs-called **Merged**.

**Table 3.**
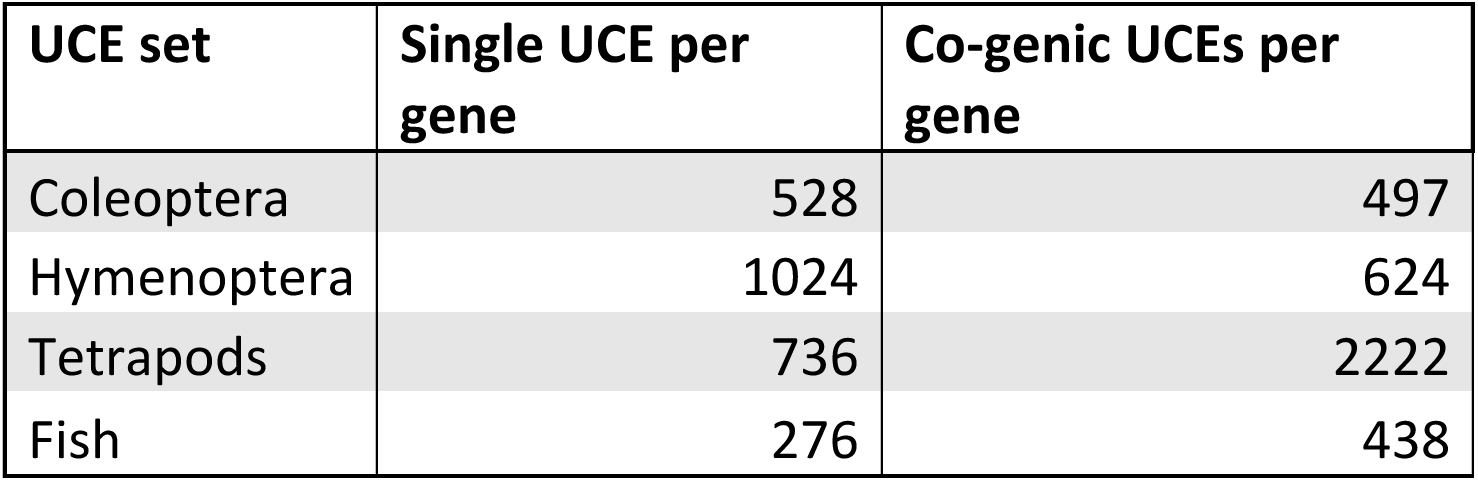
Count of the number of times single and co-genic UCEs are found in particular datasets. This count includes both exonic and intronic UCEs.

#### Calculation of distance between UCEs for Gallus gallus

The distances between co-genic UCEs (e.g. Fig. 1, see “Gene 1”) were often found to be thousands to tens of thousands of base pairs long. Merging distantly located regions of the same genes (which naturally happens in the case of transcriptome data) increases the chance of merging regions that have different recombination histories-which may impact species tree analyses (but see Lanier and Knowles 2012). To understand more about where UCEs occur in a genome in relation to each other, we used the *Gallus gallus* genome (the base genome of the tetrapod-5k-UCE probe set) and custom R script to explore the distance (in base pairs) between co-genic UCEs, as well as the distance to the nearest UCE upstream and downstream from a set of co-genic UCEs (referred to herein as nearest neighbor UCEs). For each set of co-genic UCEs, we used the results of the *blastn* analyses described above to identify where a particular UCE locus was found (position, gene and scaffold/chromosome). We then calculated the distance between co-genic. As the m8 file (results from *blastn*) is already ordered by position along a particular chromosome/scaffold, to find the nearest neighbor distances, we simply found the distance upstream and downstream from the UCEs that bookended a particular set co-genic UCEs.

### Species Tree Analyses of curated UCEs

#### Species Tree Analyses of Genic versus Intergenic UCEs

To evaluate the effects of genomic class on support and topology, we reconstructed species trees based on only **Genic** and only **Intergenic** UCEs for weevils, ants, mammals, songbirds and caragimorph fish. Using *R*/unix scrips modified from Van Dam et al. 2017, we ran a maximum likelihood (ML) analysis in *RAxML* 8.2.11 (Stamatakis 2014) on each individual UCE locus, with the “–f a” options for a rapid bootstrap analysis (100 bootstrap replicates (BS)) and searched for best scoring tree. A General Time Reversible + gamma (GTRGAMMA) site rate substitution model was used for each locus. We then constructed a **Genic** and **Intergenic** species tree for each taxon set using summary methods implemented in ASTRAL-III (Zhang et al. 2018).

#### Species Tree Analyses of Merged versus Unmerged UCEs

To evaluate the effects of merging co-genic UCEs, we reconstructed species trees based on **Unmerged** and **Merged** UCEs for weevils, ants, mammals, songbirds and caragimorph fish. For clarity, our 1) **Unmerged** UCE set is based on standard protocols and considers each UCE locus as an independent unit (one UCE locus used to reconstruct a single gene tree) and 2) our **Merged** UCE set includes co-genic UCEs merged together to generate a single gene tree, along with all remaining genic and intergenic UCEs each treated as a single locus.

For each **Unmerged** and **Merged** set of UCEs, we ran a maximum likelihood (ML) analysis in *RAxML* 8.2.11 (Stamatakis 2014) with the “–f a” options for a rapid bootstrap analysis (100 BS replicates) and search for best scoring tree. A General Time Reversible + gamma (GTRGAMMA) site rate substitution model was used for each locus. Next we constructed 2 species trees per taxon group using ASTRAL-III (Zhang et al. 2018) first using the gene trees from all **Unmerged** UCE loci and second, using the gene trees from the **Merged** dataset. We ran ASTRAL with the default settings and performed multi-locus bootstrapping (Seo, 2008).

### Effects of characterizing and curating UCEs on bootstrap values and topologies

#### ABS comparisons

To identify if there was any difference in bootstrap support between the gene trees generated from the **Genic** UCEs versus **Intergenic** UCEs and between the gene trees generated from the **Unmerged genic** UCEs versus the **Merged co-genic** UCEs, we calculated average bootstrap support values for each individual gene tree (see Supplementary Material for R and Python code). Average bootstrap support values (ABS) were calculated using a modified R script from (Borowiec et al. 2015). While non-parametric bootstrapping cannot be taken as a measure of absolute of gene tree estimation error (GTEE), it loosely correlates with it (Efron et al. 1996; Holmes 2003, 2005; Susko 2009; Molloy and Warno 2018).

#### T-tests and GLM

We performed two-sample t-tests in *R* between the means of the ABS of the **Genic** versus the means of the ABS of the **Intergenic** gene tree sets and between the means of the ABS of for **Merged co-genic** versus the means of the ABS for the **Unmerged genic** gene tree sets for all taxa to see if a UCE’s genomic category or if merging co-genic UCEs had a statistically significant effect on ABS values. We then calculated the Cohen’s d statistic in *R*. We also investigated the effect that locus length, genomic categorization, and merged or unmerged status had on ABS value using a generalized linear model (GLM).

#### Comparison of tree topologies and shapes: species trees

To assess the impact of our curation efforts on resulting tree topologies we next calculated the Robinson–Foulds distance (RF-dist) (Robinson and Foulds, 1981) tree distance metrics that rely on the tree topology in the *R* package *Phangorn* (Schliep 2011). In addition, we calculated the KF-distance (KF-dist), (Kuhner and Felsenstein 1994) which measures the sum of squares differences between individual branch lengths.

#### Comparison of tree topologies and shapes: gene trees

For assessing differences between the gene trees from the **Merged** and **Unmerged** UCE sets, we used two different tree shape metrics from their Laplacian spectrum calculated in the R package *RPANDA* (Lewitus and Morlon 2015; Morlon et al. 2016). We selected the skewness (asymmetry) of the spectral density profile and the peakedness (peak height) the largest y-axis value of the spectral density profile (Morlon et al. 2016). Normalized, each of these metrics gives a separate description of tree shape: skewness detects the relative timing between branching events (lower values indicate more branching near the stem of the tree whereas higher values indicate more branching near the tips), peakedness (lower peak height values indicate a more even tree shape, whereas higher values indicate a more ladderized tree shape).

### Assessing Species Tree Accuracy for Merged and Unmerged UCEs

To identify if the merging of co-genic UCE loci improves the accuracy of inferred species trees, we conducted a simulation study. In brief, we outline below our procedures based on slight modifications from Mirarab and Warnow, 2015.

#### Simulation methods for gene trees and species trees

As in Mirarab and Warnow, 2015 and Molloy and Warnow, 2018, we used *SimPhy* (Mallo et al. 2016) to simulate 50 species trees and their constituent 1000 gene trees, each with 50 taxa under high levels of incomplete lineage sorting (see Supplementary Material S1 for specific parameters used in simulations). We chose to simulate under high levels of incomplete lineage sorting because we expect that under these conditions, where topological inference may be at its most ambiguous, we would see performance markedly increase in for the longer “merged” loci we simulated next. Sequences were simulated under these gene trees using Indelible (Fletcher and Yang, 2009). We performed 50 replicates each with 1000 genes, the speciation rate was set to (1e-07) with a tree depth set to 1.0×10^7^ generations and a global population size of 2.0×10^5^. Sequence lengths were designated to reflect the lengths of **Merged co-genic** loci constructed from the mammal UCE dataset. Specifically, lengths were drawn from lognormal distributions with the log mean controlled by drawing uniformly between 6.2 – 10.2, which corresponds to lengths between 500 – 26,600 bp, representing the range in length of merged co-genic loci from the mammal dataset.

After creating the 50 sets of loci for each species tree, we then subdivided each of these loci to represent our **Unmerged** dataset. Here we selected loci length from a normal distribution of between 300 – 800bp, with a mean of 500 and standard deviation of 74, also representative of the mammal dataset. This resulted in 546,775 total alignments over our 50 replicates, with on average 10,936 alignments per species tree replicate.

Along with testing the accuracy of species trees based on **Merged** and **Unmerged** loci, we also tested the effects of **Randomly Merged** loci. By building species trees based on **Randomly Merged** loci, we were able to examine whether any accuracy improvements of species trees based on the **Merged** loci set were simply due to longer loci length or shared genealogy. To generate **Randomly Merged** loci, we randomly merged, without replacement, loci from the **Unmerged** dataset. The number of loci that were selected to be merged at any one time was chosen from the distribution of the number of times the **Merged** loci were divided up to generate the **Unmerged** loci set. For a distribution of the loci lengths for each of these three-simulated sets, see Supplementary Material S1.

Gene trees for our simulation study were reconstructed using RAxML under a GTRGAMMA site rate substitution model for each locus. For the species trees based on the 1000 locus datasets, we used ASTRAL-III (Zhang et al. 2018). However, for the larger **Unmerged** (∼10,000 locus) datasets we used ASTRAL-MP (Yin et al. 2019) to take advantage of AVX2-CPU and GPU processors so that the runs would finish. We then compared the resulting Robinson–Foulds tree distance between our **Merged**, **Unmerged, and Randomly Merged** species trees compared to their associated simulated “true” species trees.

## RESULTS

### Genomic Characterization of UCEs across the Tree of Life

For the 12 UCE sets examined, roughly 51%, ±16.8 95%CI, of UCEs were found within exonic regions of the genome (Fig. 2; Table 1). The percentage of UCEs located within exons varied greatly between organismal Classes; for the Insecta UCEs, 86.5% (Diptera) and 51.2% (Hemiptera) were found in exons, whereas in Vertebrata, the exon percentage varied between 41.7% (Ostariophysan fish) and 10.3% (Acanthomorph fish). The vertebrate sets tended to be composed mostly of non-coding regions compared to the invertebrates. The percentages of UCEs found in intergenic regions of the genome varied between 0.83% (Diptera) to 45.7% (Acanthomorph fish). There were some UCEs that could not be placed in an intron, exon or an intergenic region alone and were found to be spanning any two of these regions (Fig. 2). For some UCE sets, these loci were a relatively major component (see Ostariophysan fish Fig. 2), but for most UCE sets they were a small fraction or absent all together (Fig. 2; Table 1). Also, within the tetrapod UCE probe set, we found several probes that could not be recovered in more recent genome assemblies. Though this affected very few probes, it demonstrates that probe recovery may change as genomic assemblies are improved or updated, and highlights the fact that these UCE annotations are only as good as the annotations of their base genomes.

### Focal taxa UCE characterization and curation

For weevils, ants, mammals, songbirds and caragimorph fish, we filtered the UCEs located within genic regions of the genome and identified the genes that each UCE represented. We found that within Insecta, between ∼52% (weevils) and ∼62% (ants) of the UCEs are found as single UCEs in a gene (singleton UCEs) (Fig. 3). Almost as many UCEs belong to genes that are represented by more than one UCE (co-genic) (Table 1, Fig. 3). For the tetrapod UCE set, there are roughly three times as many co-genic UCEs as there are singleton UCEs. Within the caragimorph fish UCE set, ∼63% of UCEs are singletons.

**Figure 3.**
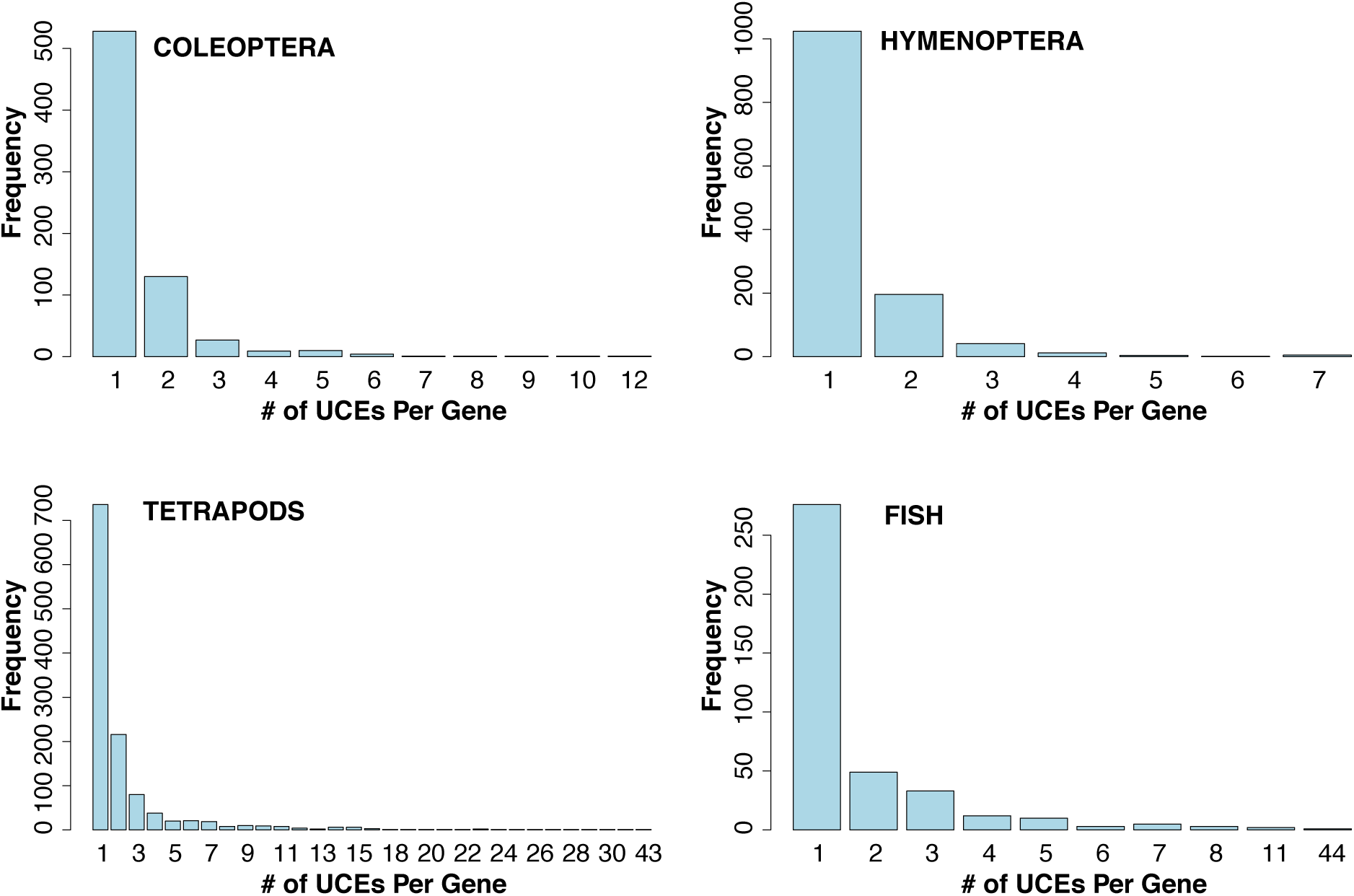
Bar-plot of frequency of the number of UCEs that can be found within a gene, e.g. ∼50 separate genes each containing 2 UCEs as seen in the lower right panel. The X-axis labels indicate the number of UCEs per gene.

#### Calculation of distance between UCEs for Gallus gallus

We found that the majority of UCEs within a gene were clumped and not widely dispersed in comparison to their nearest neighbor distances. The majority of co-genic UCEs were within 20kb to each other, with the average distance of 27kb. While some of the nearest neighbor UCEs were within 10kbp (see Fig. 4), the majority were much farther apart >100kb with the average distance of 458kb.

**Figure 4.**
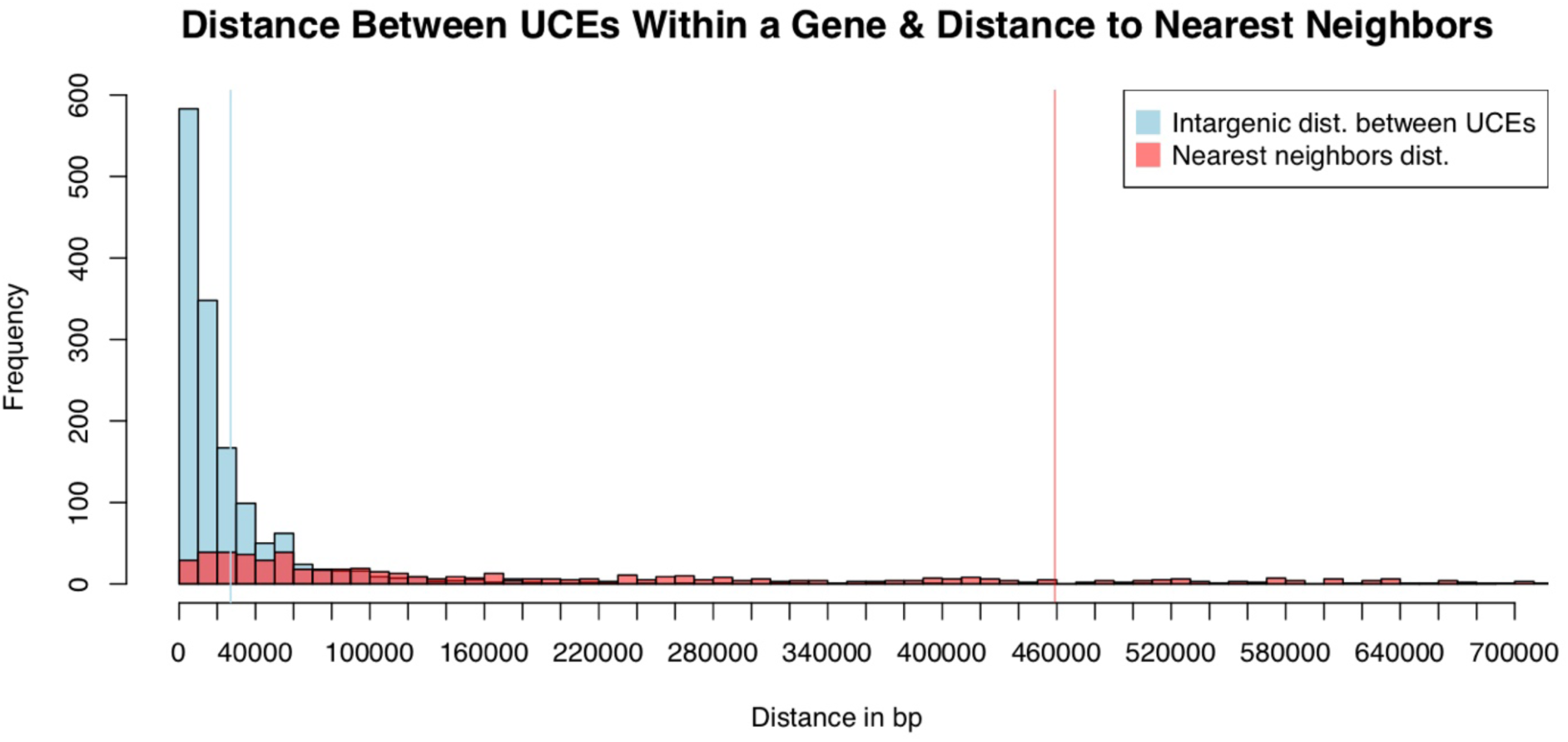
Histogram of distances between UCEs in the 5k-Tetrapod UCE set. UCEs were mapped back to the chicken genome (blastn 100% matches of probe to base over 100% of the length of UCE probe). The distances between co-genic UCEs were then measured (light blue bars), followed by measuring the distance up and downstream from a genic set of UCEs to its nearest neighboring UCE (light red bars). Vertical lines indicate the mean distance between co-genic UCEs (blue) or mean distance to nearest neighboring UCE (red) for a co-genic set of UCEs. Both estimates are local to a UCE’s scaffold/chromosome.

### Results of phylogenetic analyses of curated UCEs

#### Gene Tree Analyses of Genic versus Intergenic UCEs

Our comparison of gene trees based on unmerged **Genic** UCE loci versus those found in the **Intergenic** regions show largely no difference in average bootstrap support (ABS) (Fig. 5; Table 4). In the weevils and fish datasets, there was no significant difference between ABS of intergenic and unmerged genic gene trees, according to a t-test P-value and Cohen’s d. In the mammal set, there was significantly more support for the intergenic gene trees according to the t-test P-value but not according to Cohen’s d. Ants and birds showed significantly more support for genic gene trees (Table 4), but the Cohen’s d shows that there is a very weak difference between the ant datasets and no difference in the bird datasets.

**Figure 5.**
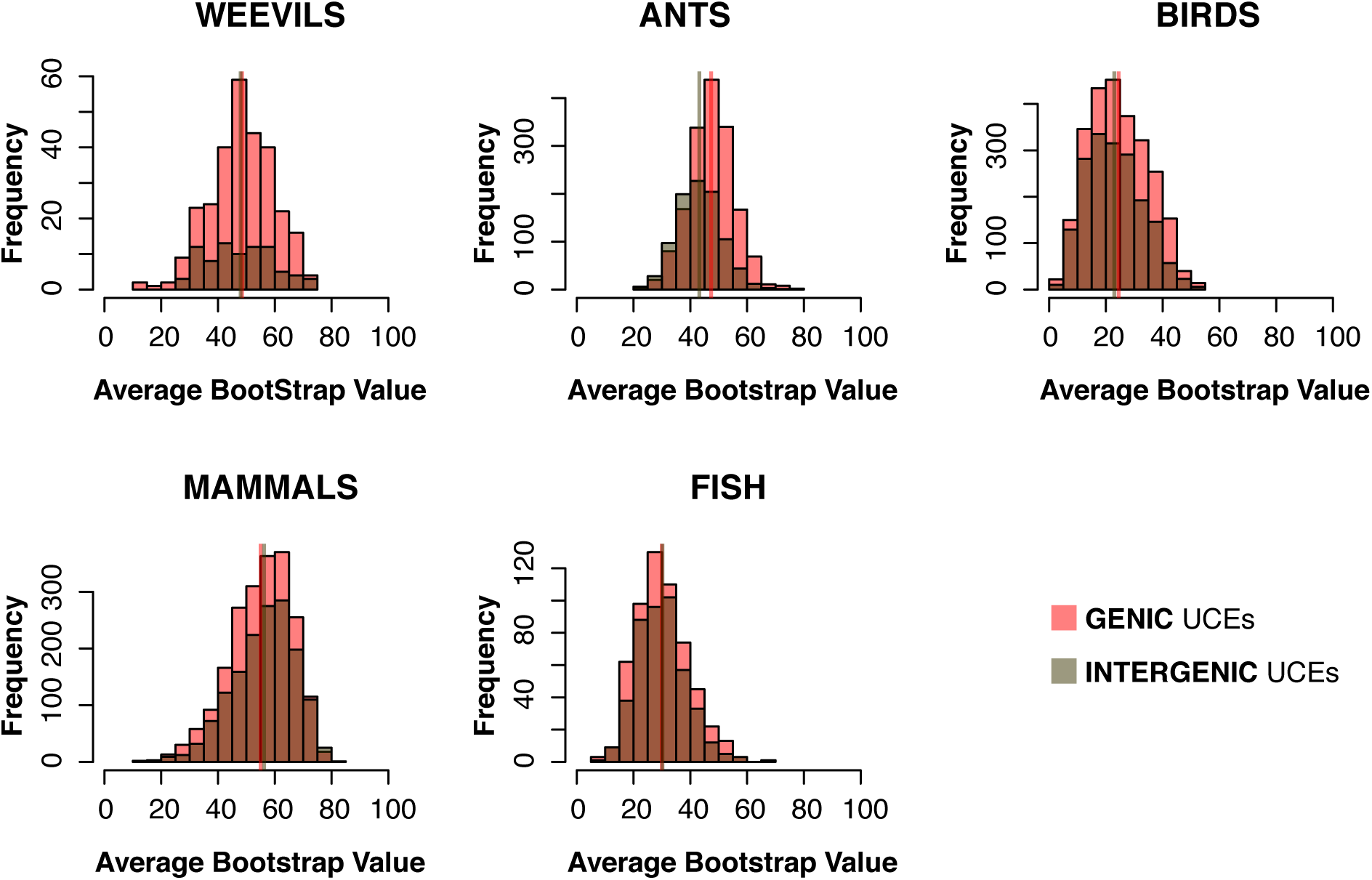
Histograms of average bootstrap support (ABS) values for gene trees based on **Intergenic** UCEs and **Genic** UCEs. Vertical lines represent the mean of ABS values for each set of gene trees (overlapping or adjacent in all but ants). Generally, the distribution of ABS values for gene trees based on **Intergenic** UCEs and **Genic** UCEs are overlapping and largely similar.

**Table 4.** Means of average bootstrap support values (ABS) per-gene tree for **Intergenic** and **Genic** UCEs across taxa. T-tests show significant differences in ants and birds, yet Cohen’s D shows no difference in ants and a weak difference in birds.

| UCE set | Mean ABS<br>INTERGENIC | Mean ABS<br>GENIC | t-test p-value | Cohen's d |
| --- | --- | --- | --- | --- |
| **weevils** | 47.96106 | 48.45831 | 0.7336 | 0.1251066 |
| **ants** | 43.17255 | 47.35368 | < 2.2e-16 | 0.2306506 |
| **mammals** | 56.19731 | 54.98488 | 0.0009687 | 0.06939678 |
| **birds** | 22.98138 | 24.49267 | 8.02E-07 | 0.03047168 |
| **fish** | 29.90741 | 30.18136 | 0.6196 | 0.04413178 |

#### Gene Tree and Species Tree Analyses of Merged UCEs versus Unmerged loci

Regarding gene tree analyses across all taxa for **Merged** versus their corresponding **Unmerged** UCEs, we found that the average bootstrap support per-gene tree was significantly higher for **Merged** UCEs than for their corresponding **Unmerged** UCEs (Table 5, Fig. 6) according to both the t-test and Cohen’s D. (Note here we are not comparing the full **Merged** and **Unmerged** datasets, but simply the merged co-genic UCEs compared to the same corresponding UCEs, yet unmerged).

**Figure 6.**
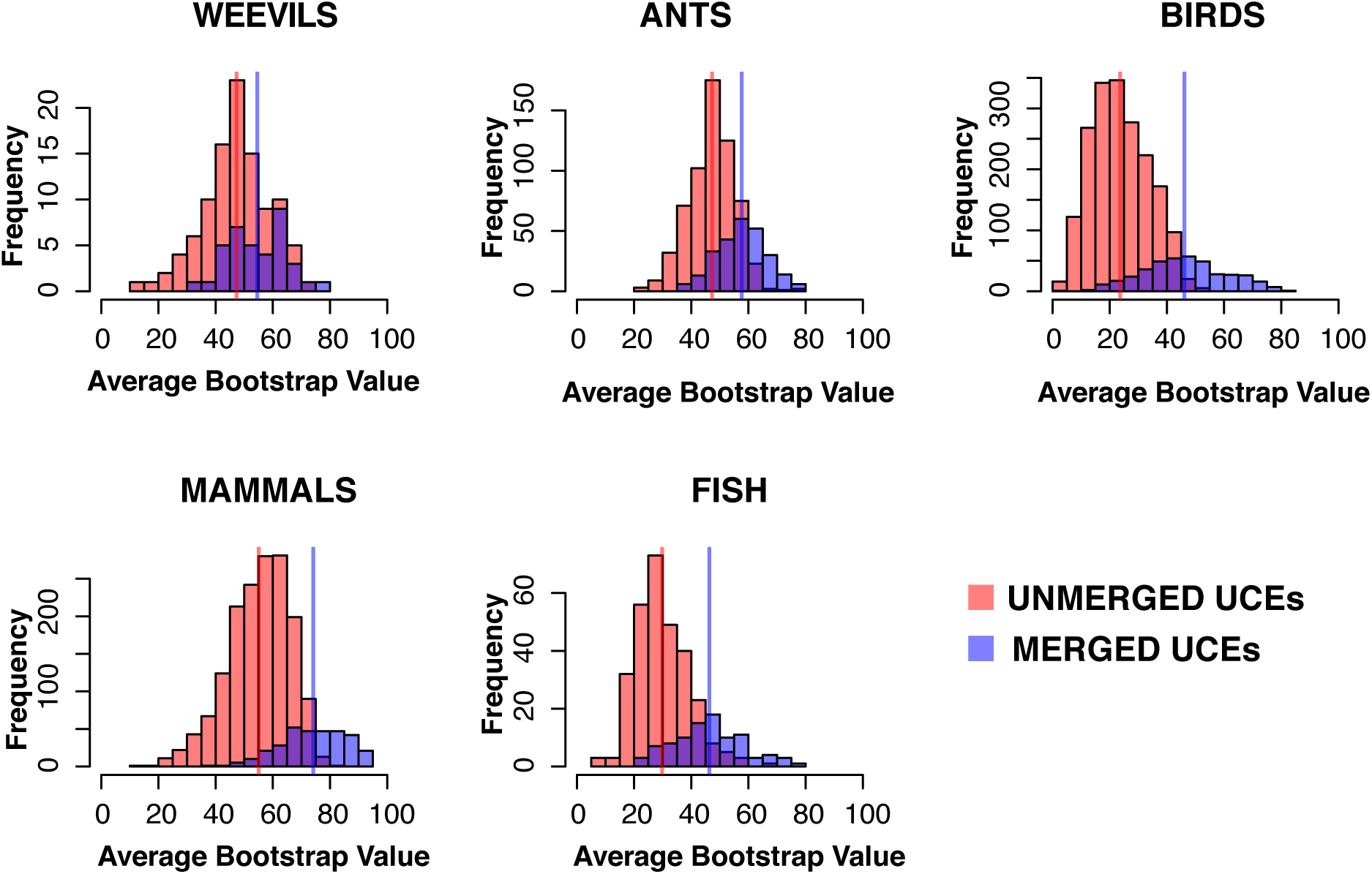
Histograms of average bootstrap support values (ABS) of gene trees generated for **Merged** UCEs and their corresponding **Unmerged** UCEs (at the exclusion of all intergenic and singleton UCEs). Red bars represent ABS values of gene trees for **Unmerged** UCEs, where each individual UCE provides a single gene tree estimate. The purple bars represent the ABS of gene trees generated by **Merged** UCEs, where all UCEs representing a particular gene were merged together to estimate a single gene tree. Vertical lines represent the mean ABS of each UCE set. The distributions are significantly different between **Unmerged** and **Merged** treatments, with the distribution of **Merged** UCE gene trees ABS values shifted towards higher support.

**Table 5.** Means of average bootstrap support (ABS) values per-gene tree and t-tests between gene trees that were merged when multiple UCEs were found within a single gene, and the same set of UCEs but unmerged and treated as single individual genes. Results show highly significant difference between **Merged** and **Unmerged** gene tree ABS values.

| UCE set | Mean ABS<br>MERGED | Mean ABS<br>UNMERGED | t-test p-value | Cohen's d |
| --- | --- | --- | --- | --- |
| weevils | 54.54468 | 47.32786 | 0.0007544 | 0.6321575 |
| ants | 57.67213 | 47.2517 | < 2.2e-16 | 1.27434 |
| mammals | 74.11935 | 55.07754 | < 2.2e-16 | 1.748992 |
| birds | 46.13401 | 23.69862 | < 2.2e-16 | 2.061103 |
| fish | 46.30876 | 29.7862 | < 2.2e-16 | 1.661292 |

The species tree results from the analyses based on **Merged** UCEs (merged co-genic UCE gene trees + all reaming UCE gene trees) and from the analyses of the standard UCE treatment (**Unmerged**, one UCE locus - one gene tree) produced similar, but not identical, results in terms of ABS and topologies. Average bootstrap support for the species tree analyses across taxa (Table 6) shows an increase in support for the trees based on **Merged** UCEs (0.29–5.13 ABS support improvement), in all but the fish species tree, which decreased by 0.38 ABS support. Average Astral quartet support values, that measure the conflict between gene trees by node, improve in **Merged** species trees across all taxa. Regions where support was weak in the analyses based on standard UCE treatment remained similar with slight improvement in the **Merged** UCEs analyses; however, these two data treatments often resulted in different topologies. Here we present an annotated topological comparison between the **Unmerged** and the **Merged** species trees for birds (Fig. 7). Comparisons of the topological differences of the remaining **Unmerged** versus **Merged** species trees can be seen in Supplementary Material S1.

**Figure 7.**
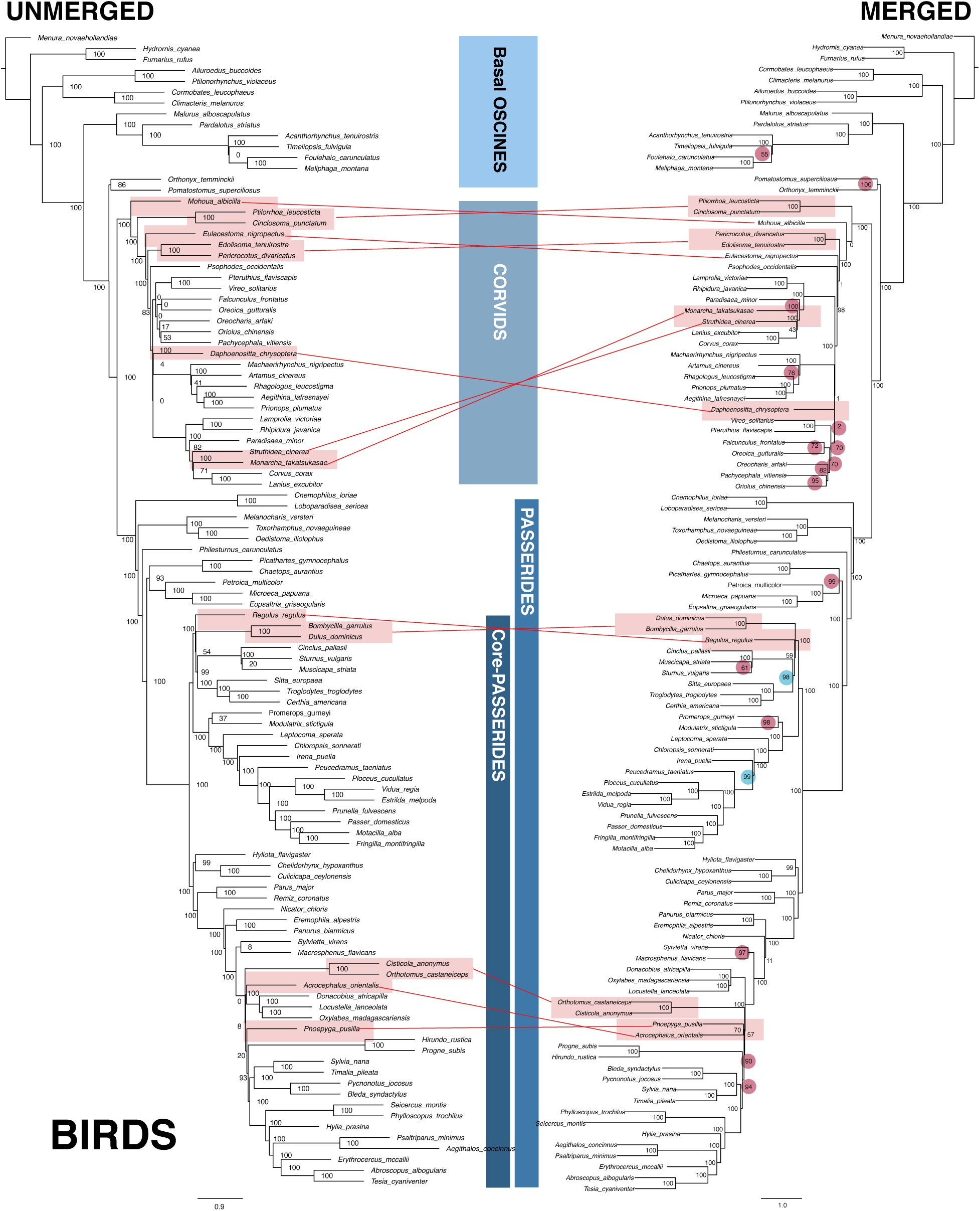
Comparison between the **Unmerged** and **Merged** species tree for birds. Differences in bootstrap values are indicated on the **Merged** tree (red circles denote increases, blue–decreases). Bootstrap values are generally higher in the **Merged** species tree and topological shifts exist within corvids and core passarides (shown with red lines).

**Table 6.** Comparison between average bootstrap support and Astral quartet support between species trees constructed in ASTRAL of Unmerged and Merged UCE sets.

| UCE set | ABS<br>UNMERGED<br>species tree | ABS<br>MERGED<br>species tree | Quartet Support<br>UNMERGED<br>species tree | Quartet Support<br>MERGED<br>species tree |
| --- | --- | --- | --- | --- |
| weevils | 90.38 | 91.11 | 58.83 | 59.84 |
| ants | 94.1 | 94.39 | 61.39 | 61.98 |
| mammals | 93.65 | 96.23 | 69.29 | 71.05 |
| birds | 86.1 | 91.23 | 50.56 | 52.65 |
| fish | 89.73 | 89.35 | 55.40 | 56.62 |

#### GLM

We found a general trend that ABS values were highly correlated with locus length (Table 7; Fig. 8). Another influential factor contributing to model fit of the GLM was the merging of co-genic UCEs in all but the weevil dataset. In contrast, including unmerged UCEs as a category did not significantly improve model fit in all but the ant dataset, which showed slight improvement.

**Table 7.**
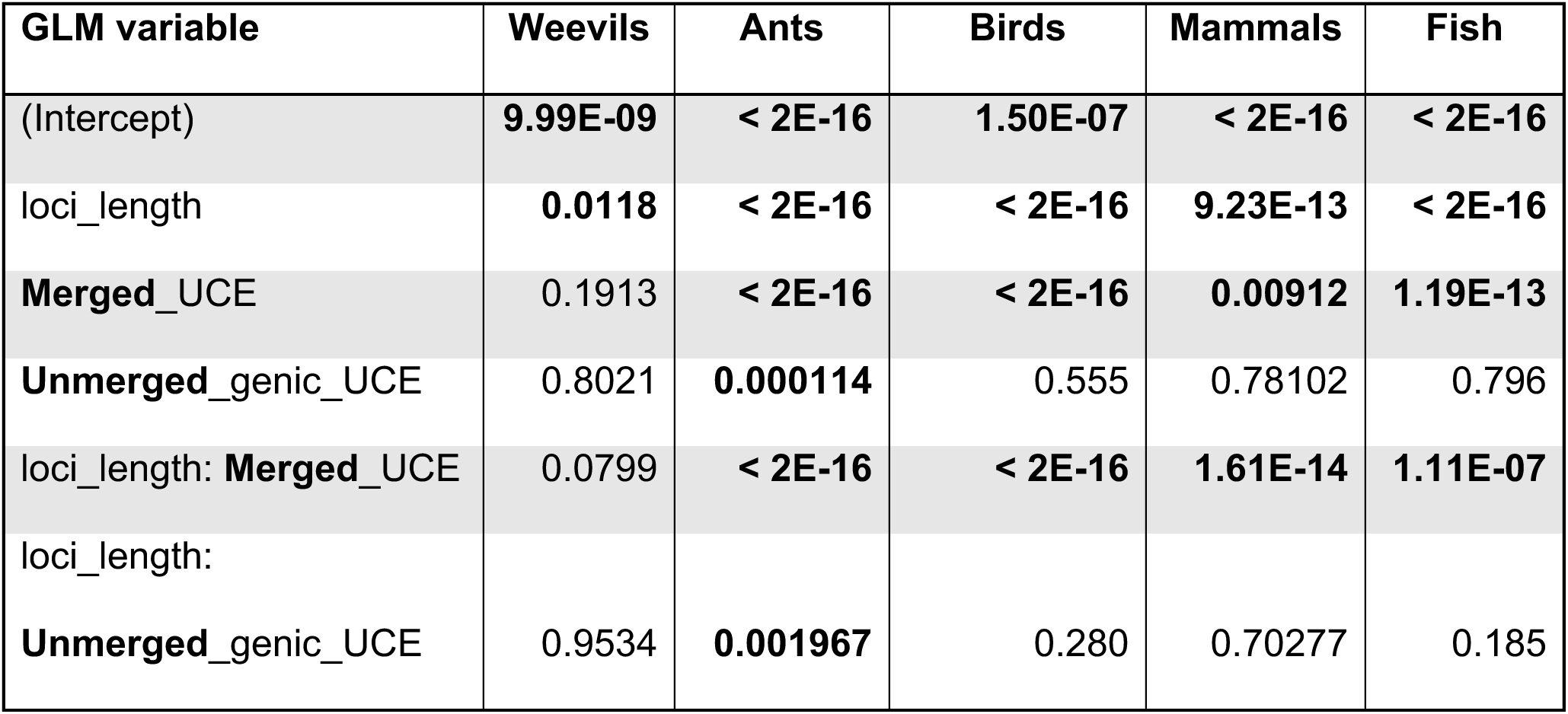
P-Values from GLM analyses. Loci length was consistently found to be a significant influence on average bootstrap support across datasets. Merging of UCEs also strongly influenced loci length and thus had an effect on average bootstrap support. Significant P-values (less than 0.05) are highlighted in bold font.

**Figure 8.**
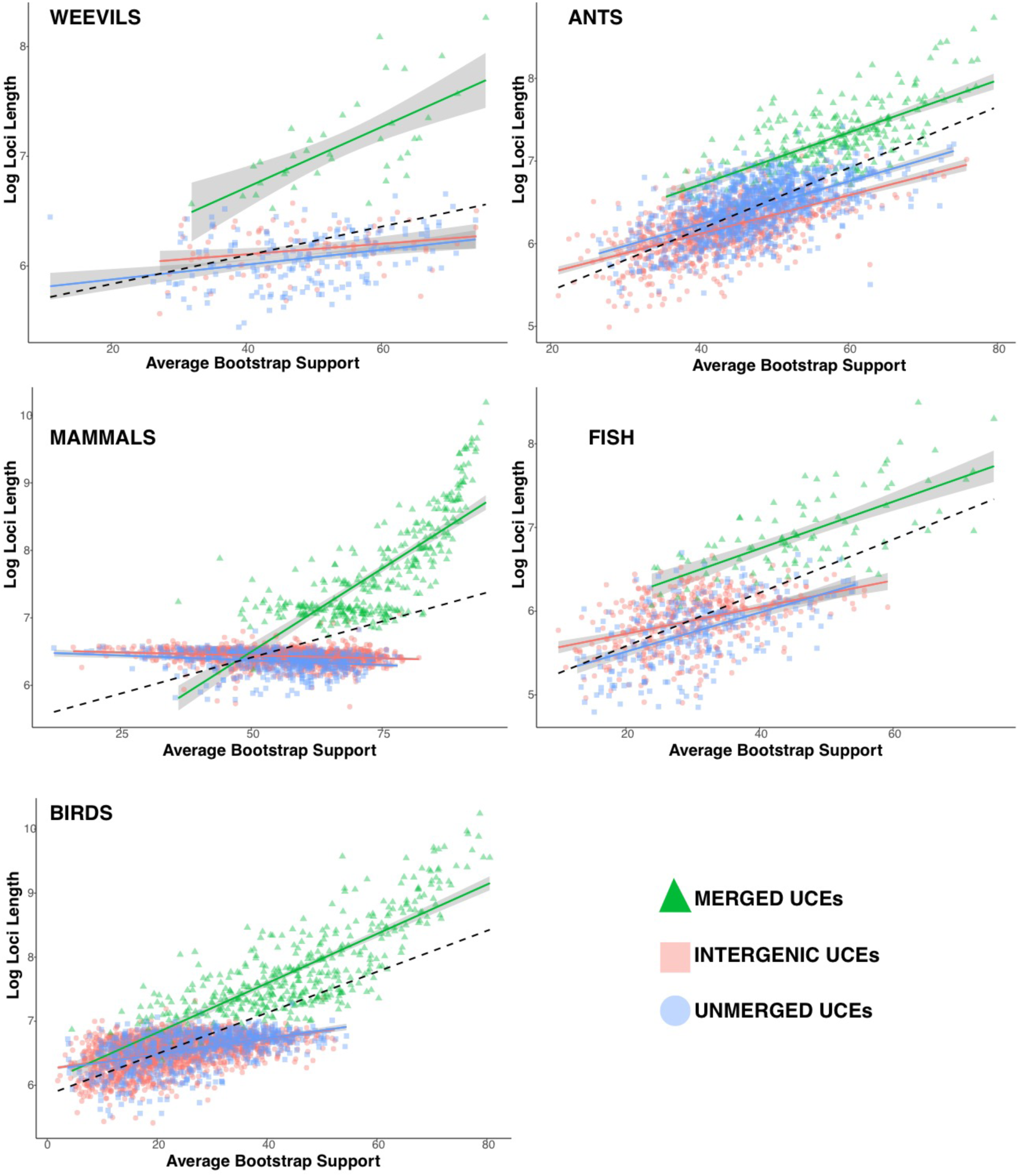
Linear regression of log of UCE length verses average bootstrap support (ABS) values of its corresponding gene tree. Categories include **Intergenic** (all UCES that fell in intergenic regions), **Merged** (all merged co-genic UCEs), and unmerged (all remaining genic UCEs that were single representatives of single genes). Dashed line represents the correlation of ABS versus log of loci length for all gene trees. Gray regions around regression lines represent 95% confidence interval. ABS values generally increase with increasing loci length.

#### Comparison of tree metrics

The RF-dist distances and KF-dist distances were computed for the **Genic** versus **Intergenic** species trees and the **Merged** versus **Unmerged** species trees for all taxa (Supplementary Material S1). For **Genic** versus **Intergenic**, RF-dist ranged from 6 (mammals) to 22 (ants), and KF-dist distances ranged from 1.3 (mammals) to 4.4 (ants). For **Merged** versus **Unmerged** species trees, RF distances ranged from 0 (ants) to 20 (birds) and KF-dist ranged from 0.52 (fish) to 1.63 (birds).

The results from the spectral density profiles, including skewness and peakedness (peak height) measures of gene tree shape, both show that on average the distribution **Merged** trees shape is less variable and more similar to one another, whereas their component **Unmerged** gene trees are more widely dispersed (see Fig. 9). In addition, **Merged** versus their component **Unmerged** species trees occupy subtly different regions of trees shape–with most significantly shifted into another region (Supplementary Material S1). These metrics largely indicate that the **Merged** gene trees are converging on a narrower and slightly different region of tree shape. The spectral density profiles for the component **Unmerged** and **Merged** gene trees for mammals are shown here (Fig. 9), while the remaining taxa can be found in the supplement (Supplementary Material S1).

**Figure 9.**
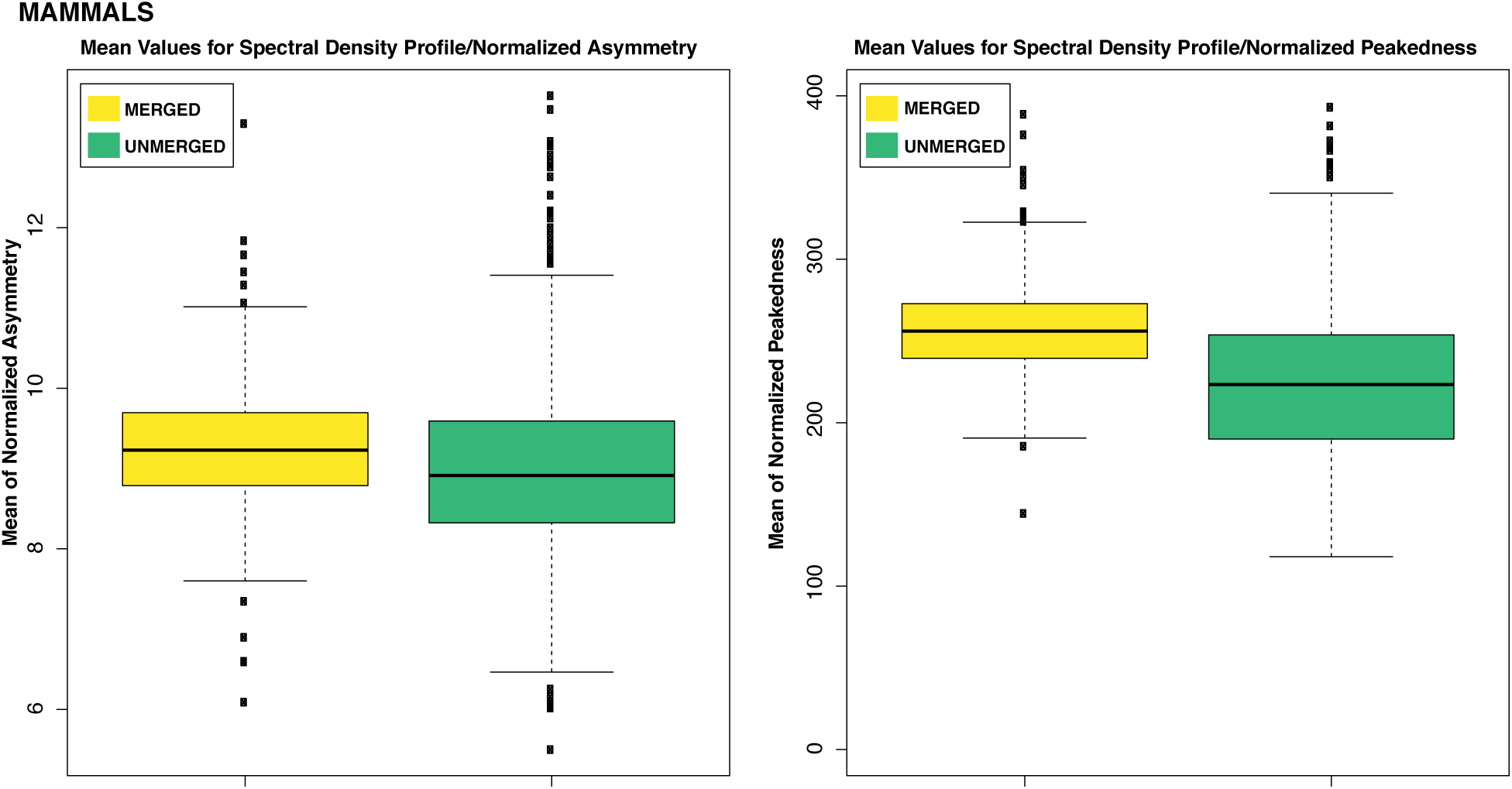
Box plots of normalized Laplacian spectral density profiles, skewness (left panel) and peakedness (right panel) for both **Merged** (yellow) and their component **Unmerged** (green) gene trees from the mammal data set. Both tree shape metrics show that the **Merged** gene trees have a narrower range of values. In addition, their means are slightly shifted into a different region of tree space compared to their component **Unmerged** gene tree spectral density profile metrics.

### Assessing Species Tree Accuracy for Merged, Unmerged and Randomly Merged loci

The results of the simulations clearly favored the **Merged** loci (which are essential just long loci) over loci that were **Unmerged** (the long loci broken up). The **Merged** gene trees reconstructed the original (true) simulated species tree in 84% of the simulations, the **Randomly Merged** 74% of the time, whereas the **Unmerged** gene trees found the original species tree only 2% of the time (Fig. 10). The average RF-dist between the true species tree and the **Merged** species tree set was the smallest (mean RF-dist: 0.4, SD 0.9), slightly higher between the true species tree and the **Randomly Merged** species tree set (mean RF-dist: 0.8, SD 1.8), and greatest between the true and **Unmerged** species trees (mean RF-dist: 88, SD 18).

**Figure 10.**
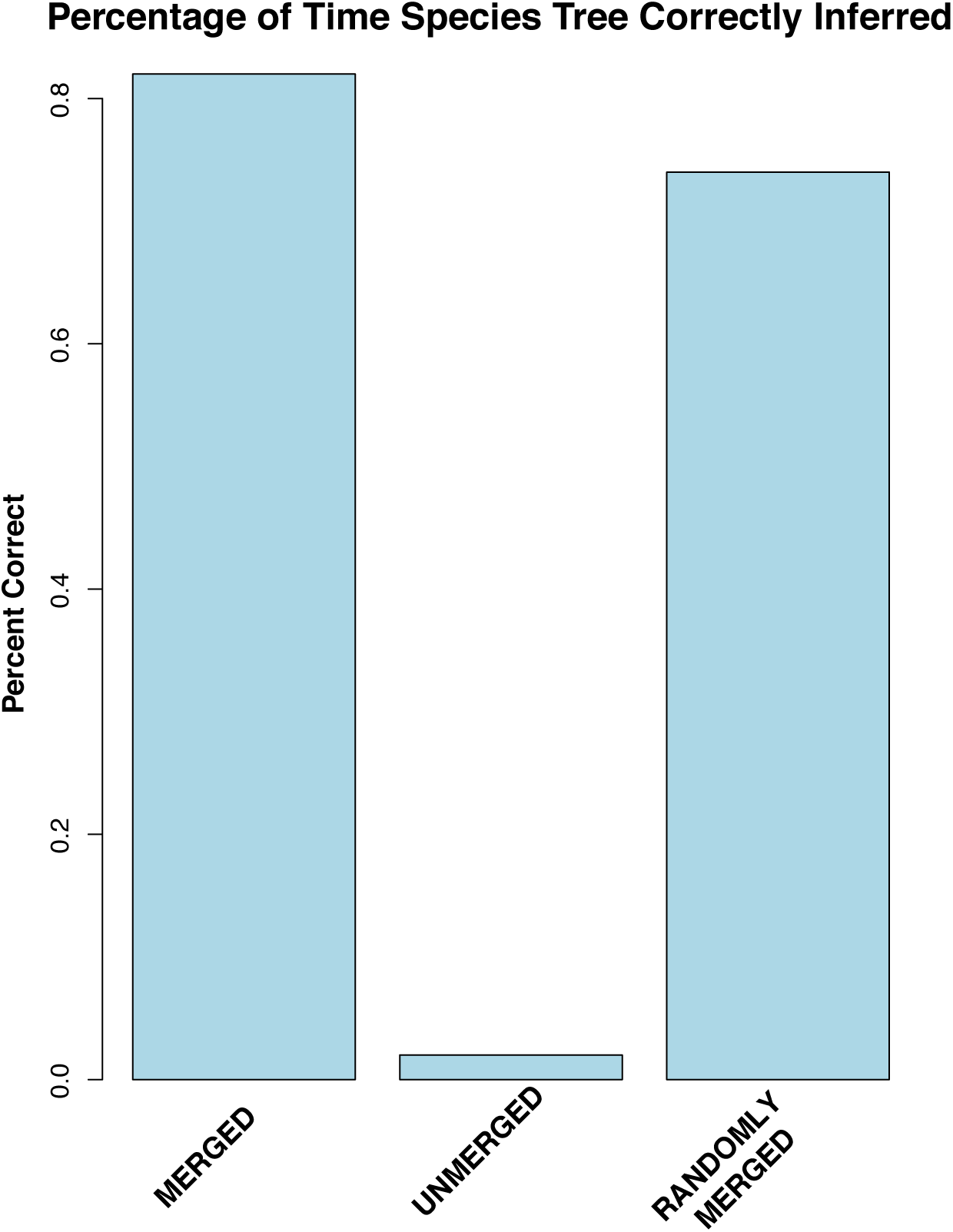
Robinson and Foulds tree distance between the true species tree and the results of the **Merged** (long simulated loci), **Unmerged** (the long loci broken up) and **Randomly Merged** loci (the broken-up parts of the original long loci randomly merged together). The average RF-distance for the inferred tree to true tree are: 2.2 **Merged**, 90.6 **Unmerged**, 3.1 **Randomly Merged**.

## DISCUSSION

Here we characterized UCE sets from a diversity of organisms and found that in each UCE set there are loci from exonic, intronic and intergenic regions. The identification of coding regions in UCEs has been mentioned in previous studies, but not thoroughly explored (Branstetter et al., 2017b, Jarvis et al. 2014). Interestingly, the genomic identity of UCEs appears to vary between invertebrates and vertebrates, with invertebrate UCEs being primarily coding and vertebrate UCEs being mostly non-coding (Fig. 2). It is important to note that our characterizations were based on recent annotations of the base genomes (or assigned base genome) for each UCE set (see Supplementary Material S1). Our categorization of UCEs as exonic relied on gene annotations based on transcriptomes and, in many cases, algorithmic predictions. Thus, we expect that over time some of these UCE characterizations will change in accordance with updated annotations of the base genomes. Also, the annotations for the base genomes for these UCE sets are in varying stages of completeness, yet in most cases more complete than the genomes for other taxa included in these UCE studies. Because we assume orthology of UCEs between taxa within a single study, we expect that the genomic categorization of these UCEs for the base taxon also extends to the UCEs from other taxa (though equally complete and annotated genomes would be required to test this further).

UCEs were first described from the mouse and human genomes as non-coding regions (Dermitzakis et al. 2003), and so this characterization appears to have then been carried over to all organisms, though it is perhaps only partially true for vertebrates (Fig. 2). An explanation for the contrast between the genomic characterization of vertebrate versus invertebrate UCEs is unclear. The taxa that the invertebrate and vertebrate UCE sets were designed across share common ancestry at a similar age of ∼300 million years ago (Bethoux 2009; Smith and Marcot 2015), so the difference is not necessarily driven by the evolutionary age of the lineage. It is possible that the contrasting breakdown of invertebrate and vertebrate UCEs instead relates to the quality of the assembled genomes used in probe design, variation in the probe design pipeline used for vertebrate and invertebrate UCE sets (Faircloth et al. 2012, Faircloth 2017), or more interestingly, genome size and evolution.

Our results on the categorization of vertebrate UCEs are similar in composition to those reported from UCEs identified between mouse and human genomes (McCole et al. 2018). It has been suggested that UCEs play a role in genome stability, as they are enriched in contact domains (McCole et al. 2018), and have been shown to also exhibit elevated synteny (Dimitrieva and Bucher, 2012). In mice and human genomes, boundary regions flanking contact domains, as well as loop anchors, are relatively depleted of UCEs; however, the UCEs that do occur in these regions are disproportionally exonic and play a suggested role in splicing (McCole et al. 2018). It is possible that invertebrate UCEs are more often pulled from boundary regions flanking contact domains and loop anchors to explain their high exonic content.

Genomic categorization is increasingly relevant in phylogenetic analyses. Studies based on coding regions often yield varying results based on how genes are analyzed, either as amino acids or by nucleotides or by only certain nucleotide positions, sometimes depending on the age of the radiation being assessed. UCEs have been treated as non-coding units even when combined with exons, but see (Jarvis et al. 2014; Bossert et al. 2018). Yet, once coding regions from UCEs are characterized, subsequent analyses can take into consideration results based on amino acids versus nucleotides and specific nucleotide positions.

Additionally, the phylogenetic utility of varying genomic regions has been considered by authors, particularly in respect to bird phylogeny (Jarvis et al 2014; Edwards et al. 2017). Non-coding regions’ potential for species tree reconstruction has been highlighted over coding regions because non-coding regions (particularly CNEEs, Conserved Non-Exonic Elements) have less possibility of GC bias and differences in evolutionary rate, as well as an ease of alignment. When we consider the distribution of ABS values from gene trees reconstructed from **Intergenic** versus **Genic** UCEs, we found no general pattern differentiating them (Fig. 5; Table 4). The genic and intergenic gene trees result in a similar, overlapping distributions of bootstrap values across taxa.

When we considered genic UCEs, we not only found that many were exonic, but we also found that genic UCEs often occur in multiples within a single gene or are co-genic. Across taxa, co-genic UCEs were more numerous than UCEs that were single representative of particular genes, except in the Hymenoptera UCE set. Although many UCEs are co-genic, they have been historically treated as independent loci in species tree analyses, thus over-representing particular gene trees in the summary species tree analysis. By merging together all co-genic UCEs, we ameliorate the issue of non-independence (at least in these obvious cases).

After analyzing the gene trees of these merged loci, we found that the ABS of **Merged** gene trees are significantly higher than these same loci **Unmerged** (treated as singletons) (Fig. 6; Table 5). **Merged** and **Unmerged** gene tree topologies varied as well, with the distribution of **Merged** gene tree topologies generally showing less variability and covering a narrower region of tree space according to spectral analyses that consider branch length and tree shape (evenness vs ladderized) (Fig 9, Supplementary Material S1). This suggests that the longer merged loci provide more decisive phylogenetic signal. Also, the ABS of the **Merged** species trees were improved across taxa (excluding the fish dataset, Table 6) and topologies varied from those based on standard protocols (Fig. 7, Supplementary Material S1).

In general, we find that loci length was the predominate driver for the increased ABS based on merging UCEs by gene (Fig. 8; Table 7). Loci length has been found to be highly correlated with phylogenetic information content (Faircloth et al. 2012; Portik et al. 2016; Branstetter et al. 2017a; Edwards et al. 2017; Van Dam et al. 2017). Given the relationship demonstrated between ABS and accuracy of estimated gene trees (Liu et al. 2015; Molloy and Warnow 2018; Zhang et al. 2018), we expected that, broadly speaking, the merging of UCEs from the same genes into longer loci would result in more highly supported gene trees that would improve the accuracy of the resulting species tree. Our simulation study supported this hypothesis. Our investigation of the accuracy between simulated sets of **Merged** (complete, long loci), **Unmerged** (subdivided loci, the length of standard UCEs) and **Randomly Merged** UCEs, decisively favored the merged loci in terms of their ability to recover the correct species tree under high levels of incomplete lineage sorting. This finding indicates that it is not only loci length that contributes to enhancing bootstrap support values and improving the accuracy of tree inference, it is also the shared phylogenetic signal in the **Merged** loci. **Randomly Merged** loci, however, did outperform **Unmerged** loci, again pointing to the importance of loci length even at the expense of shared phylogenetic history across a locus (Adams and Castoe 2019; Bayzid and Warnow, 2013). Our simulations suggest that using fewer longer loci is preferable to many shorter, less informative loci.

Our results are also in accord with other studies (Edwards et al. 2017; Adams and Castoe 2019) that identify that longer loci with more information are preferable to shorter less informative ones, despite the fact that longer loci increase the probability of spanning recombination blocks. The effect of recombination on summary species tree methods has received recent attention (Lanier and Knowles 2012; Gatesy and Springer 2014; Edwards et al. 2016; Jennings 2017). A series of papers by Gatesy and Springer suggest that recombination misleads species tree methods (Gatesy and Springer 2014, Gatesy and Springer, 2018) and thus the authors advocate for concatenation methods. Yet a simulation study by Lanier and Knowles, 2012, found recombination to not have an overtly negative influence on coalescent-based phylogenetic analyses under high levels of incomplete lineage sorting (though only relatively short loci were considered- and see Gatesy and Springer, 2018).

Another advocated approach to address the issue of recombination in species tree analyses has been to select loci that are separated by an intrachromosomal distance threshold (Jennings 2017) to satisfy the evolutionary independence assumption of coalescent-based phylogenetic methods (Arbogast et al. 2002). In some UCE studies, UCEs within 10k base pairs of each other were discarded to avoid physical linkage (Faircloth et al. 2013; Alfaro et al. 2018)-a physical recombination distance estimated for fish. However, the accurate estimation of recombination blocks across diverse non-model organisms is currently an unrealistic approach for phylogenomics, considering that recombination rates even within species show substantial heterogeneity across the genomes of individuals (Comeron et al 2012).

Here we did find, as an example, that within the *Gallus gallus* genome, co-genic UCEs were generally clustered together within a distance of 20kb, while the distance between co-genic UCEs and their next neighbor outside of the gene were much longer (>400 kb on average). This clustering suggests that merging multiple sections of the same gene may not be as problematic in regard to chromosomal distance as suggested by Springer and Gatesy (2018). Though 20k base pairs may be longer than some suggested distances (Drosophila 12.5 kb: Jennings 2017), it is also shorter than some (Tiger salamander 17 kb-1.7 Mb: Jennings 2017). Again, this highlights the ambiguity surrounding the determination of appropriate recombination distances and where they stop and start over potentially millions of years of evolution.

In addition, the “merging” of distinct and perhaps distant regions of a single gene naturally occurs in the production of a transcriptome, and coalescent-based phylogenetic analyses based on transcriptome data are widely used (Lin et al. 2014; Fernández et al. 2018; Esstelystyn et al., 2017; Wipfler et al., 2018). The combined effect of linkage, recombination, and selection on species tree accuracy still remains unclear. In general, the interplay between recombination and selection (e.g. selective sweeps and recombination hotspots) has not been adequately addressed in the systematics debate between recombination and its effect on the accuracy of multispecies coalescent methods. Yet, as more fully syntenic genomes become available (Dudchenko et al. 2017) this more nuanced investigation can begin.

When multiple UCEs fall within a single gene, treating them as independent units in species tree analyses potentially over-represents a single gene in species tree analyses (but see Scornavacca and Galtier, 2017). The perimeter of this evolutionary unit is dependent on how genes are defined. Here we used the GFF files from well-annotated base genomes that had predefined units termed genes. This process of identifying genes in a genome by default necessitates the importance of high-quality genome annotation and the criteria/methods used in the annotation (e.g. identifying genes from robust transcriptomes and or algorithmically). The chicken genome, for example, was annotated through masking repetitive regions and then using transcripts (cDNA and ESTs) and RNA-Seq data to identify the potential genes, which were then filtered further using standard gene (codon) models (Warren et al. 2016). This high-quality annotation lessens human arbitration in gene definition, as defining genes is primarily based on biological evidence.

UCEs are increasingly important and frequently used in phylogenomics due to their accessibility in specimens of varying quality, relatively low cost, as well as due to the existence of a user-friendly bioinformatics pipeline (Faircloth, et al. 2102). Amongst genomic subsampling methods, though, they also return the shortest loci on average (Karin et al. 2019). The capture of short loci is a benefit when dealing with degraded DNA from old specimens, but short loci are a disadvantage in multispecies coalescent-based phylogenetic analyses. Short loci can also contribute to gene tree estimation error, which in turn hampers species tree inference. When using species tree methods based on UCE data, merging co-genic UCEs may help reduce the negative impacts of uninformative loci and result in a more highly supported and potentially more accurate phylogenetic estimate.

## Acknowledgments

The authors would like to thank Brian Simison and Joe Russack (California Academy of Sciences, CAS) for access to AWS servers. We would also like to thank the CAS Center for Comparative Genomics Reading Group for discussion of this paper. Thanks to the Doolin Foundation for Biodiversity and NSF 1754376, 1856402 for funding this research. We would also like to thank Adam Leaché, Brant Faircloth and the other 3 reviewers for their useful comments that improved the quality of this manuscript.

## Supplementary Material

UCE Identification Pipeline: *R* and *Python* code: https://github.com/matthewhvandam/integrating-functional-genomics-into-phylogenomics

Supplementary Material S1: tree comparison metrics and reference tables: link pending submission:

