## Supplementary Material Appendix S1 for "Genomic characterization and curation of UCEs improves species tree reconstruction": READ_ME Supplementary Material Appendix S1.pdf

This folder contains the following:

1.) Folder containing Box-Plots of gene tree spectral distances comparisons (“GENE TREE SPECTRAL DISTANCES”). These are between the unmerged and merged gene trees. There are two plots for each taxon, the peakedness and asymmetry.
2.) Folder containing the tree files for all the species trees (“SPECIES TREES - unmerged/merged - genic/intergenic”). They are broken down into two subfolders one for unmerged and merged species trees and the other for the genic and intergenic species trees, support values are bootstrap support. Trees with the quartet support values at the nodes are in the “Quartet Support” subfolders.
3.) A PDF labeled “Comparison of unmerged to merged species trees”. This displays the species trees generated in ASTRAL-III, unmerged ASTRAL tree is to the left, merged genic loci plus all remaining loci to the right. Node values indicate bootstrap support values. Changes in topology are circled on the merged species tree. Increases in ABS values are in red, decreases are in blue.
4.) Histogram of locus lengths for simulated gene tree data sets (“UCE_SIMMS_LOCI_LENGTH_with_mean.pdf”). Vertical lines indicate the mean for each simulated set.
5.) A table for the simulation parameters used (“Simulation Parameters.xlsx”).
6.) A table with the pairwise comparison of species tree distances, Robinson–Foulds distance (RF) and Kuhner–Felsenstein (KF).
7.) A table listing the UCE set and the corresponding NCBI reference numbers for the base taxon.
