## Supplementary figures and images for "Genomic characterization and curation of UCEs improves species tree reconstruction"

### 3 Comparison of unmerged to merged species trees.pdf

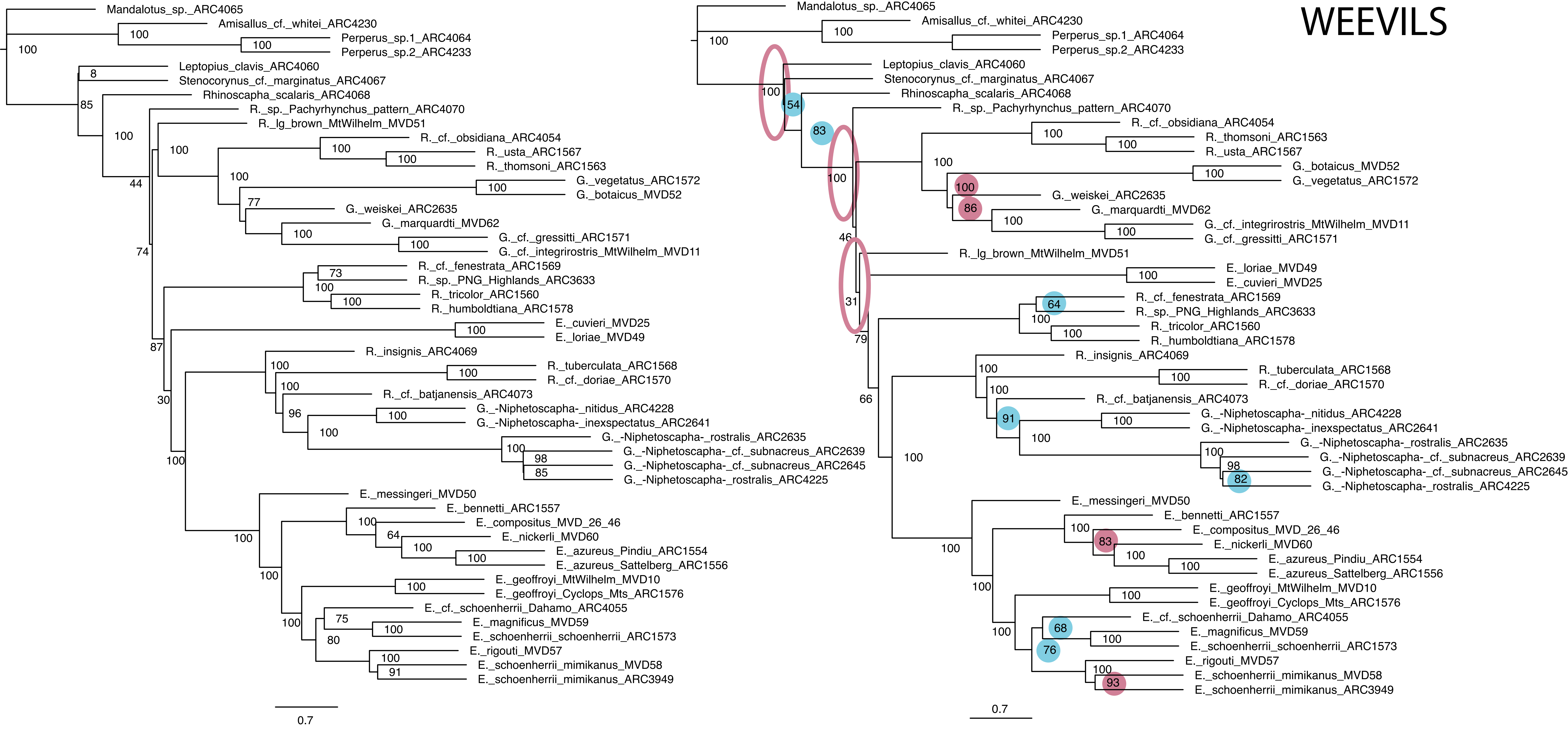

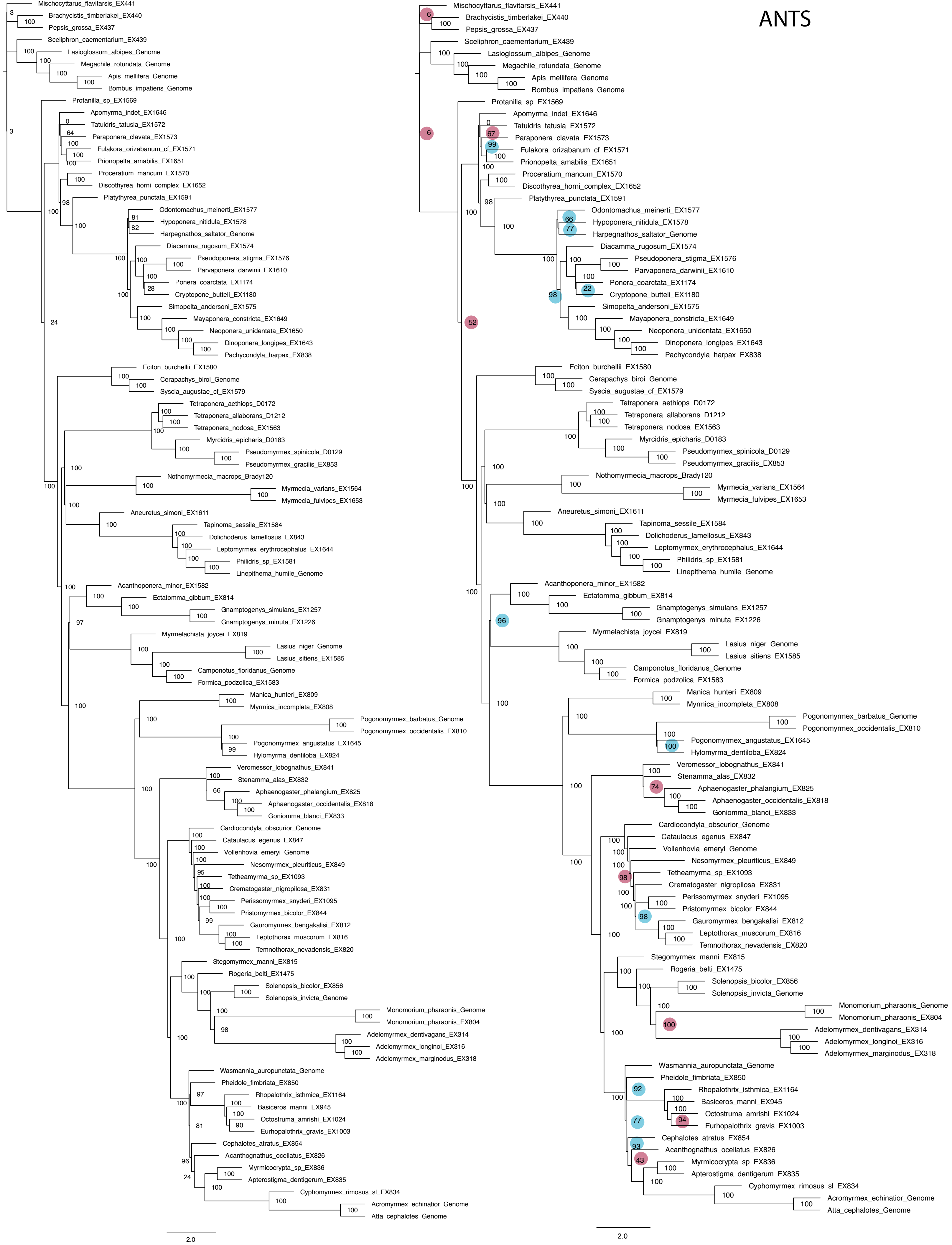

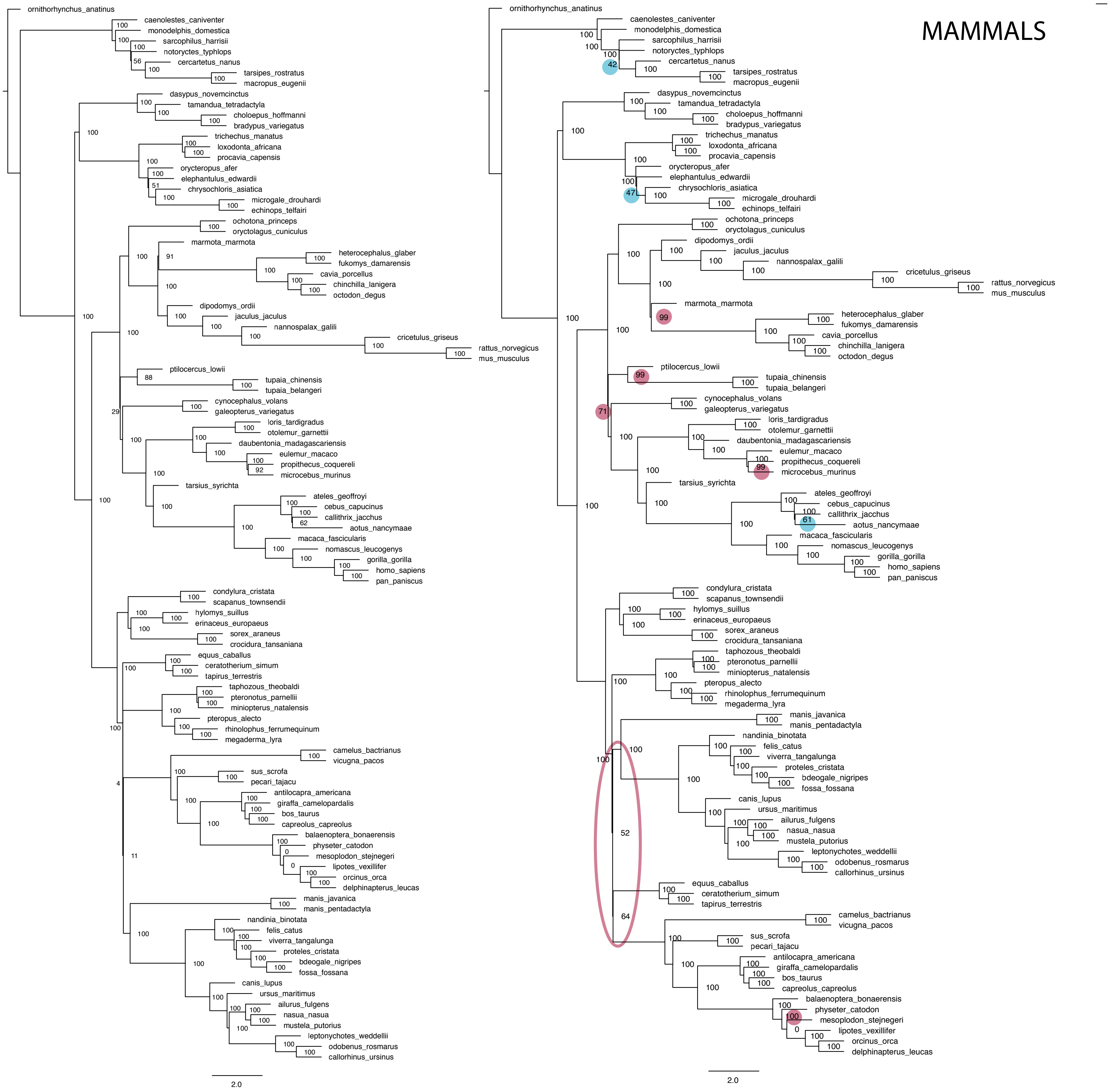

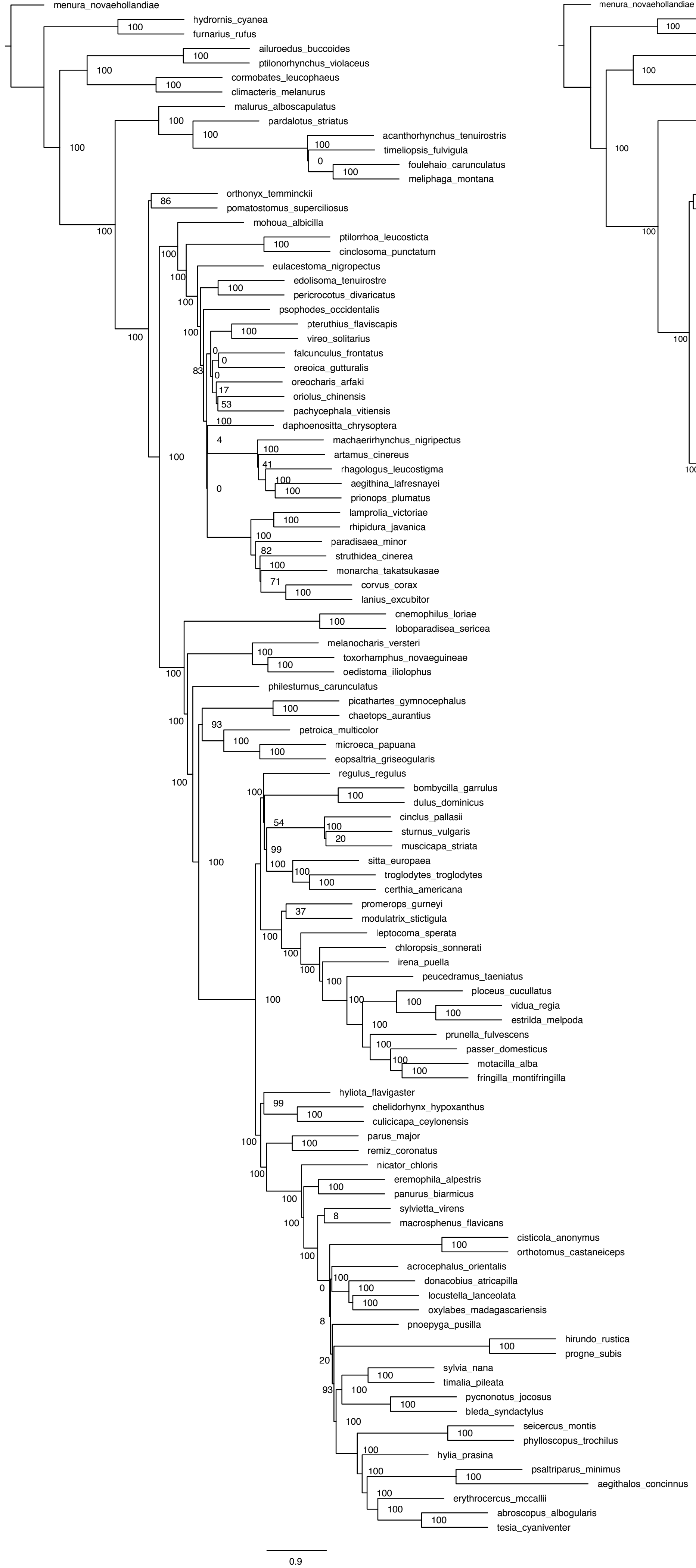

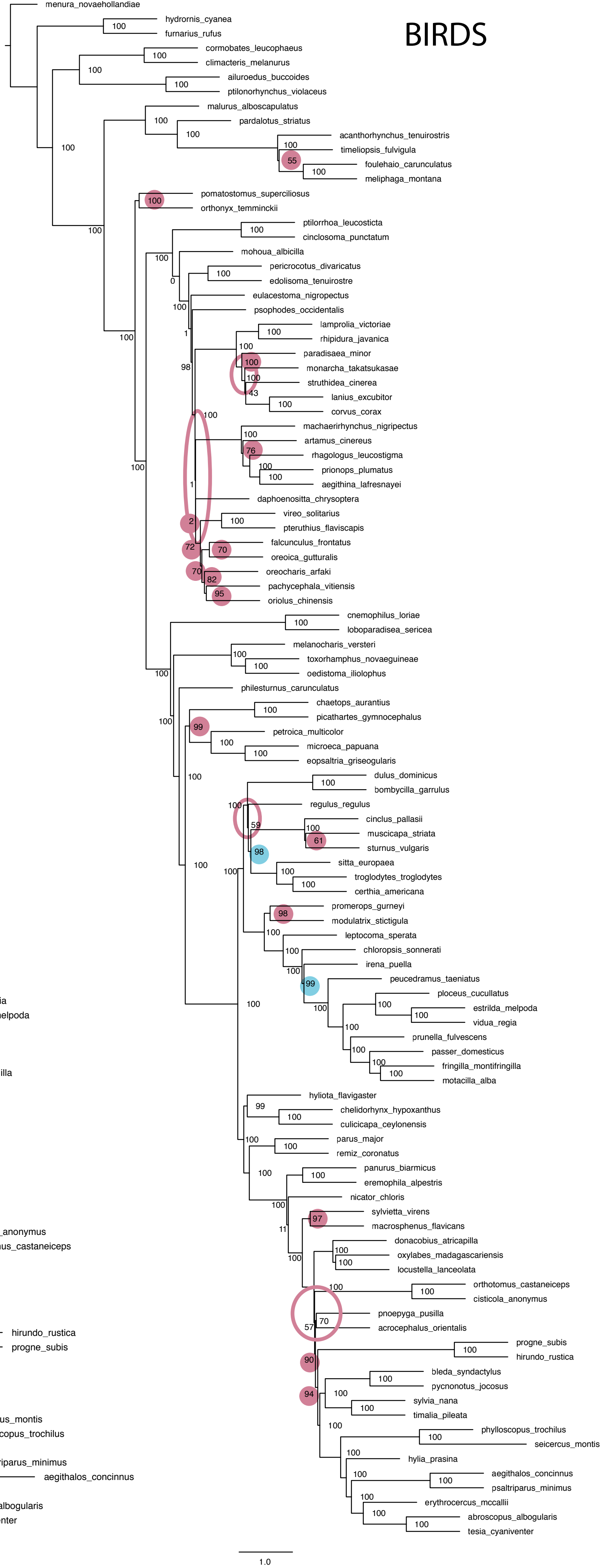

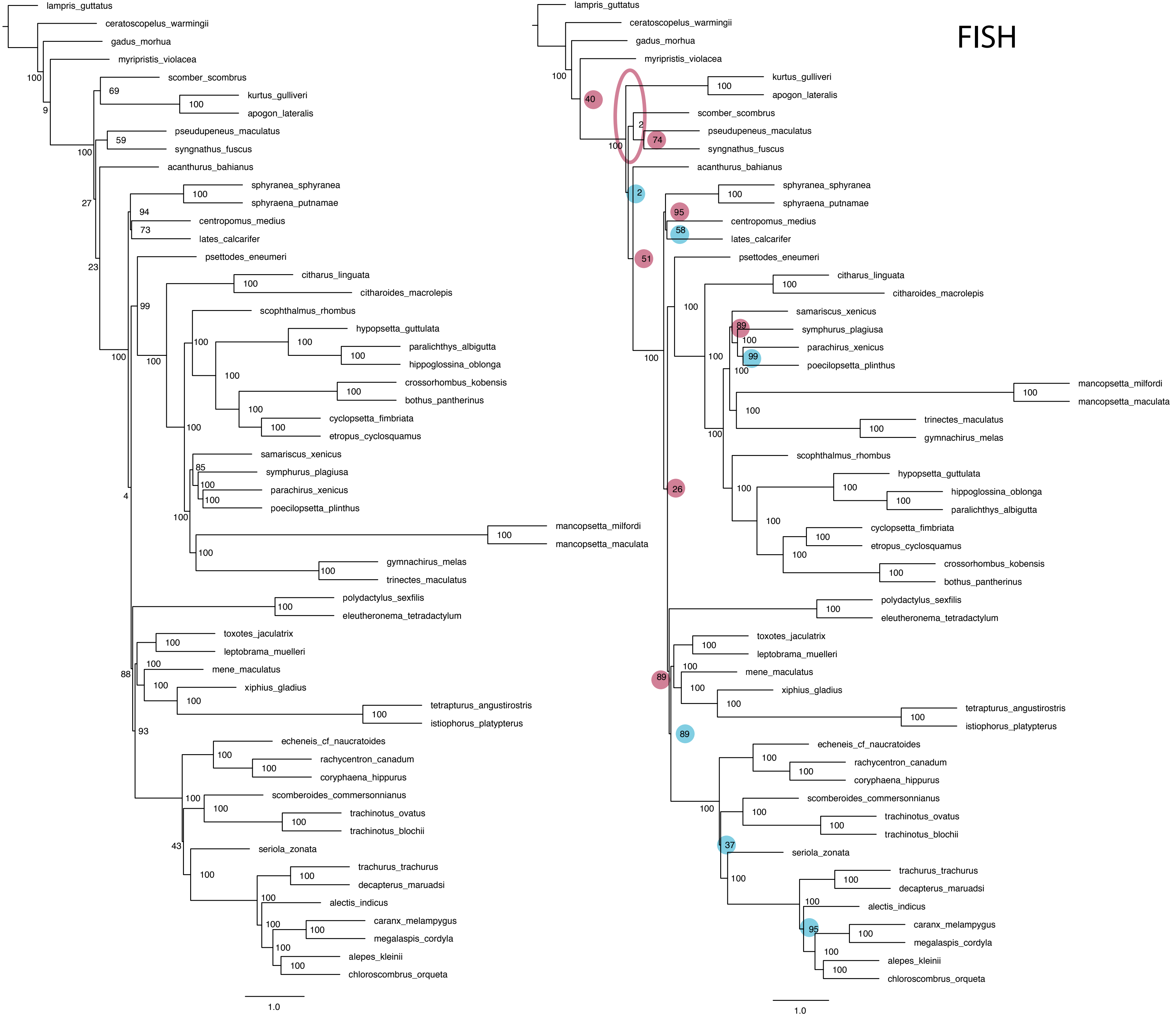

### 4 UCE_SIMMS_LOCI_LENGTH_with_mean.pdf

#### Locus Length of Simulated Gene Trees

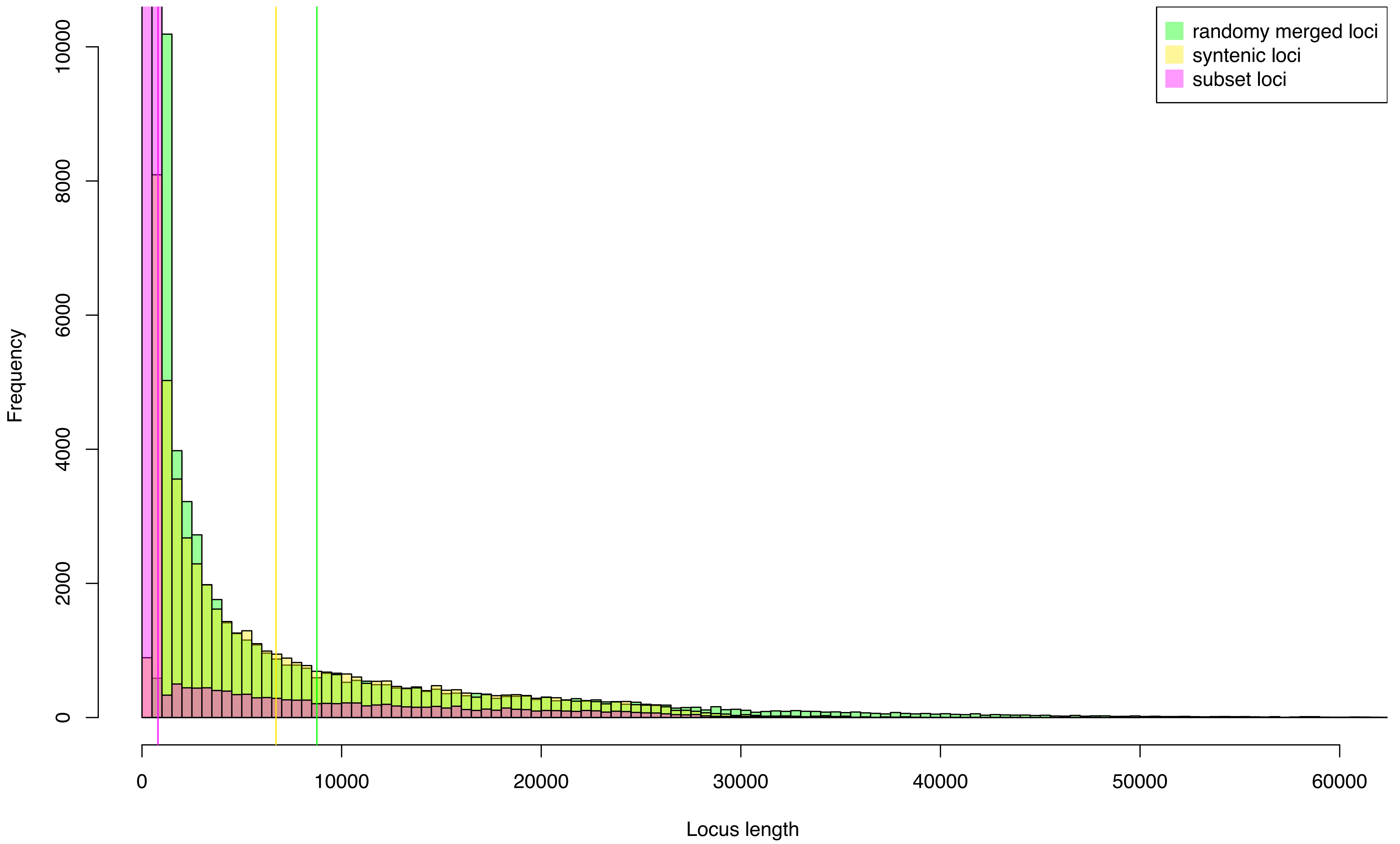

### Spectral_Density_Profile_Normalized_Asymmetry-Birds.pdf

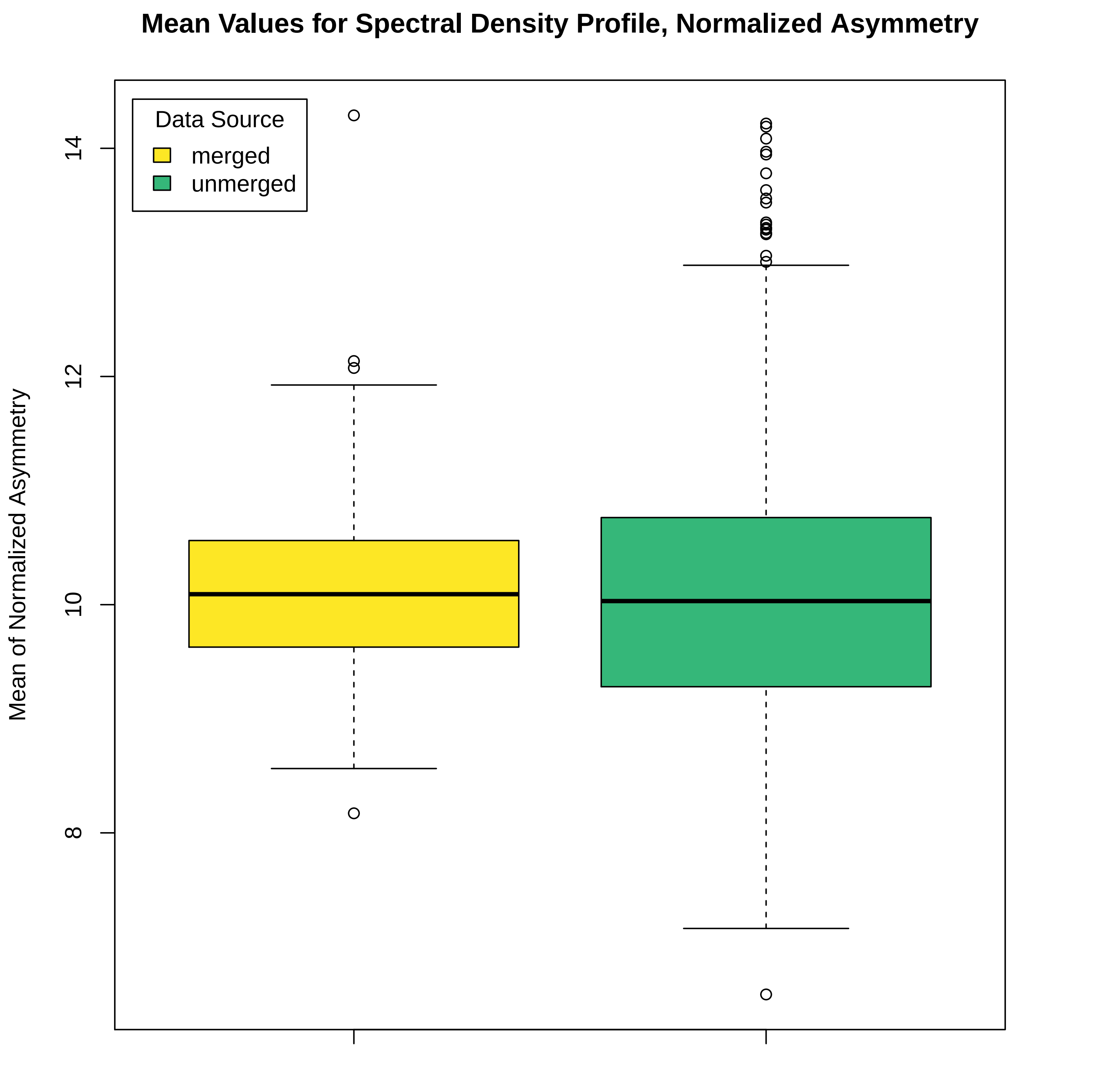

### Spectral_Density_Profile_Normalized_Asymmetry_Ants.pdf

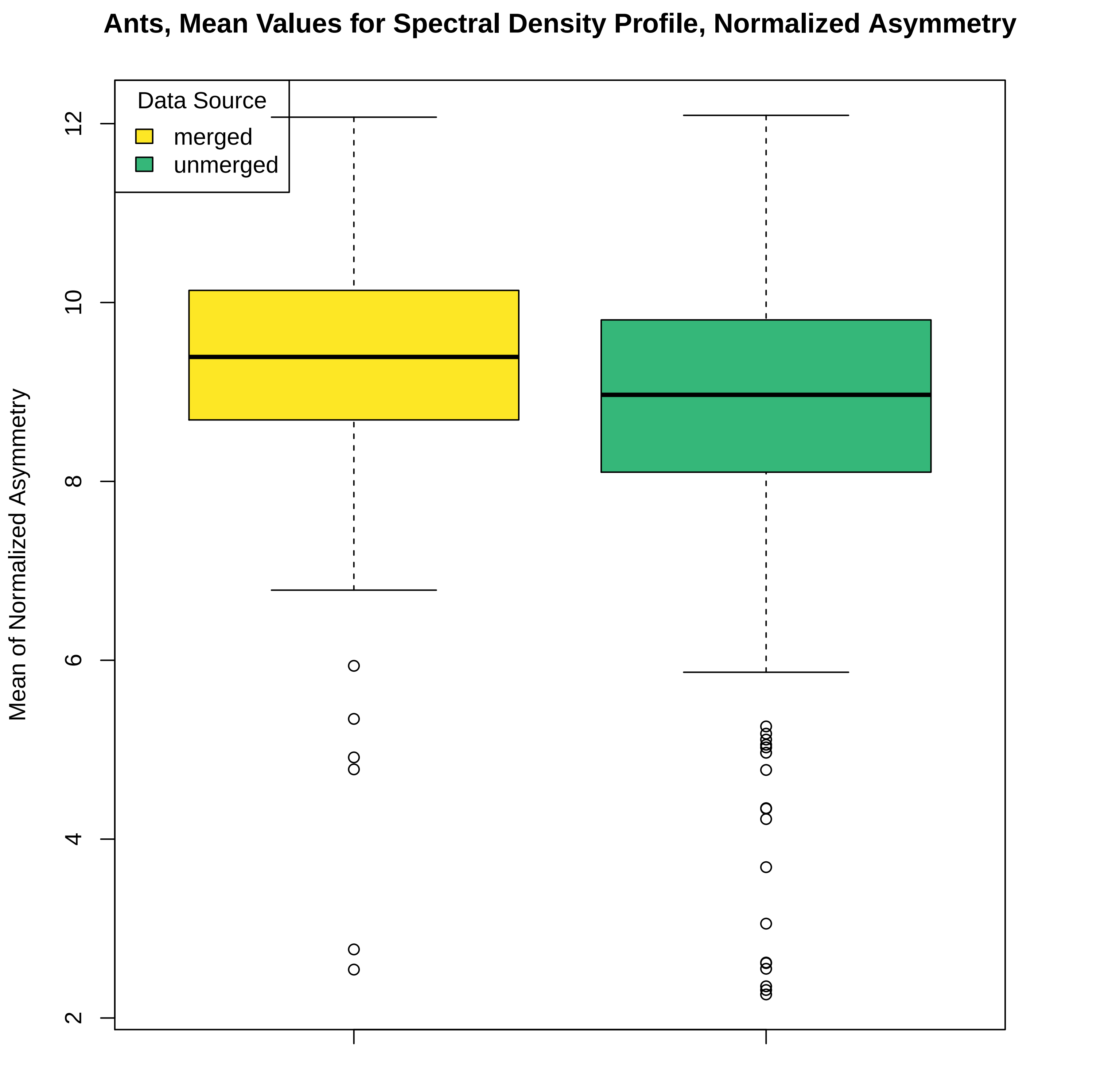

### Spectral_Density_Profile_Normalized_Asymmetry_FISH.pdf

##### Fish, Mean Values for Spectral Density Profile, Normalized Asymmetry

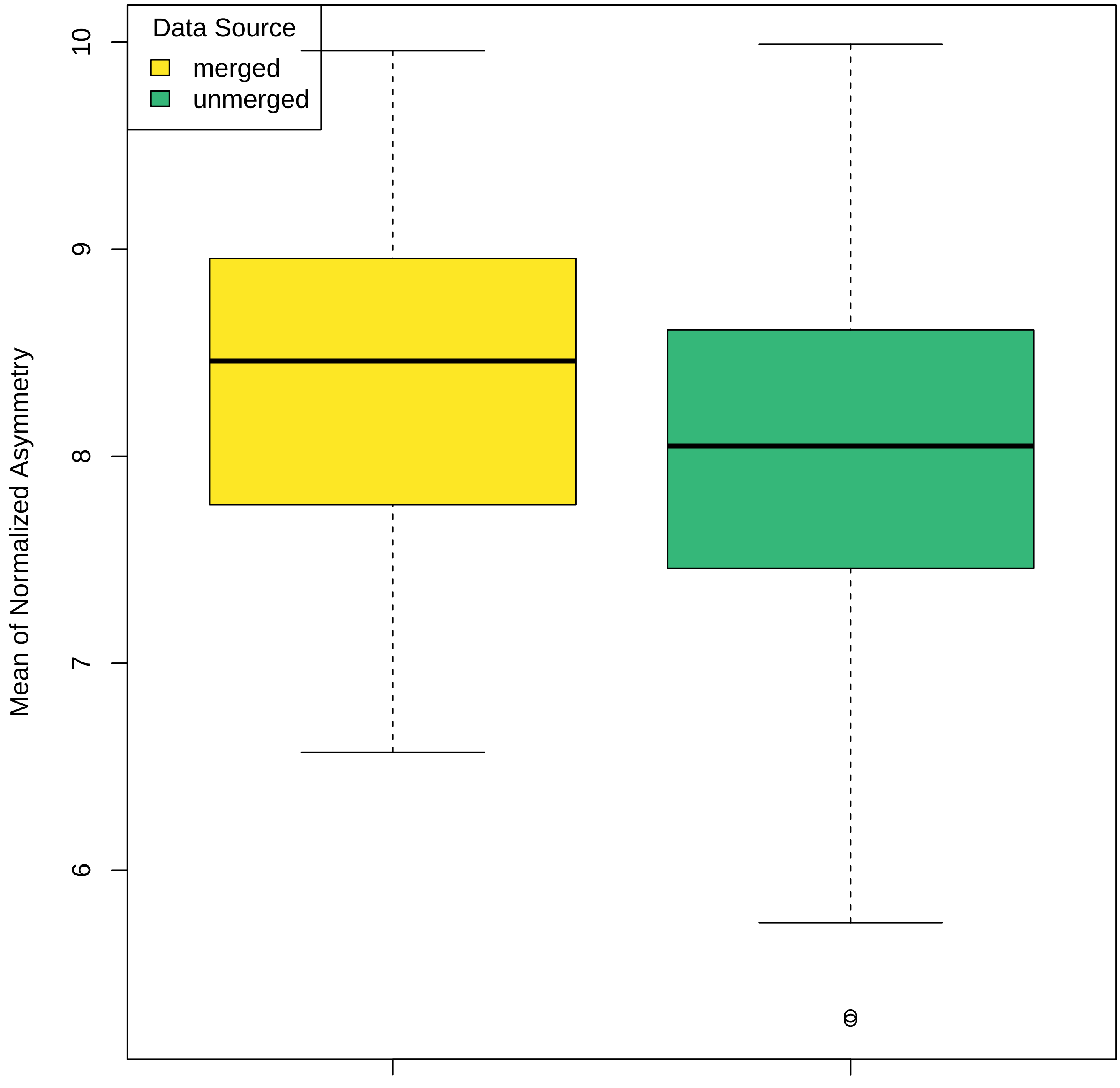

### Spectral_Density_Profile_Normalized_Asymmetry_Mammals.pdf

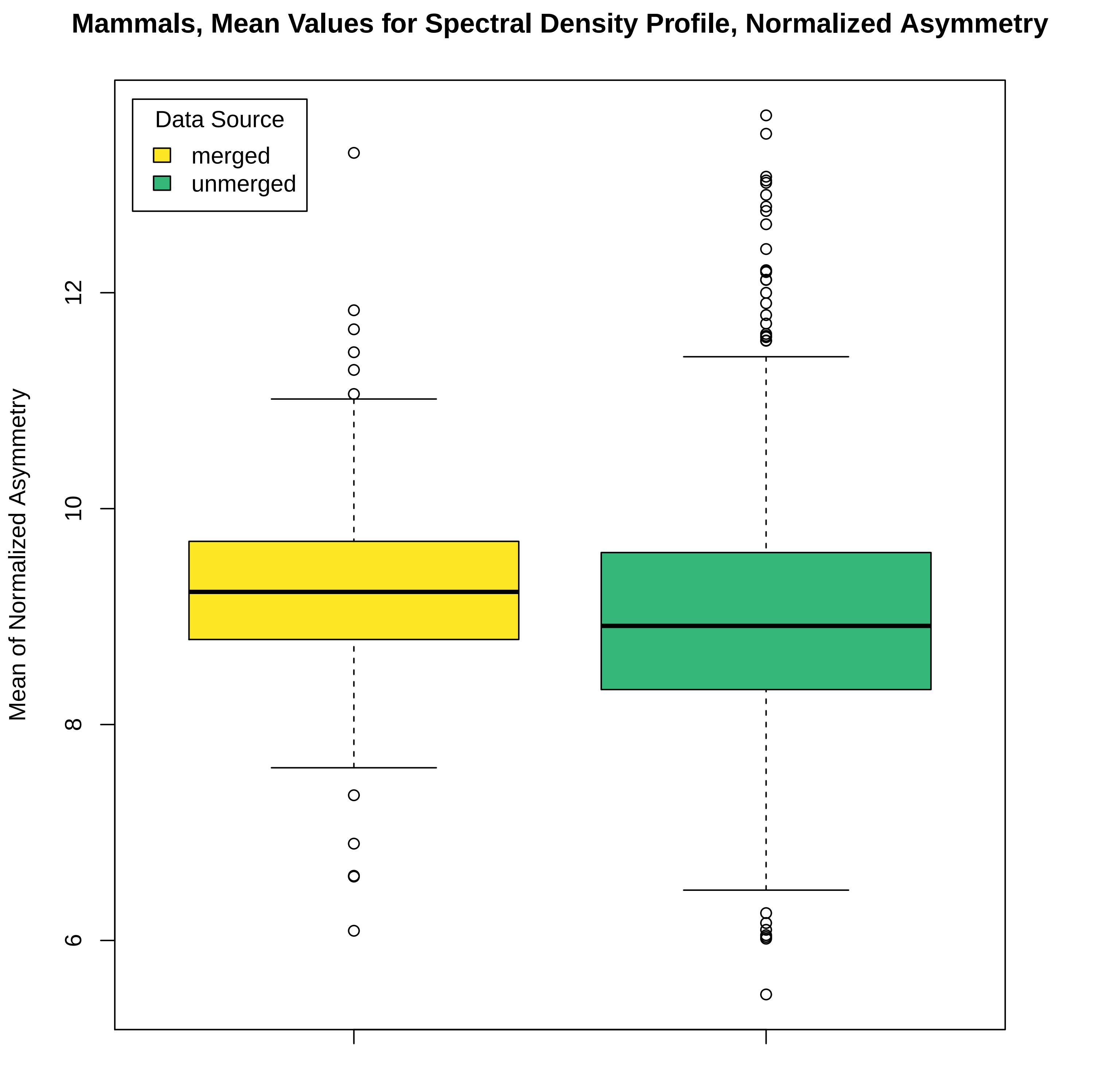

### Spectral_Density_Profile_Normalized_Asymmetry_Weevils.pdf

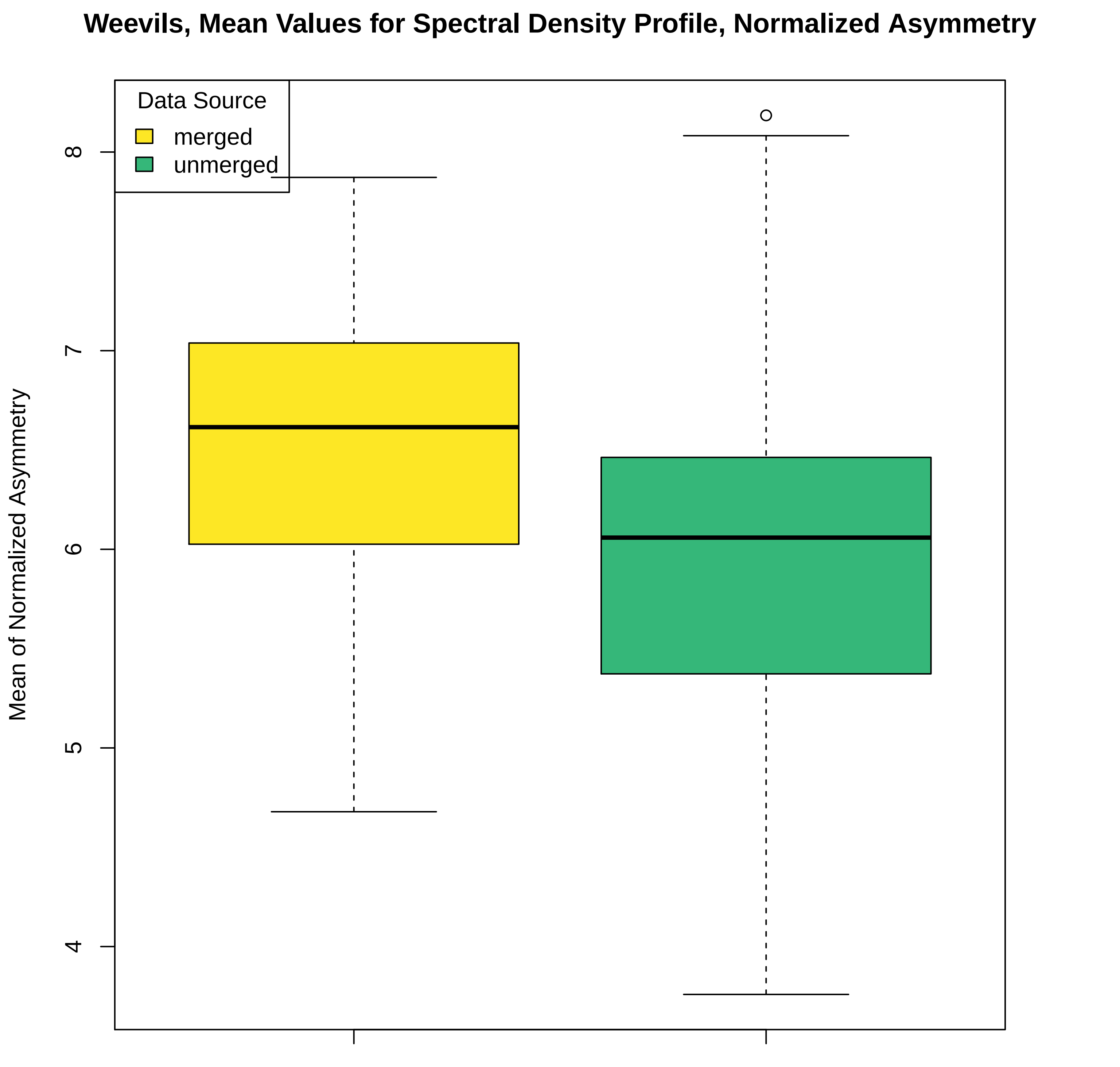

### Spectral_Density_Profile_Normalized_Peakedness_Ants.pdf

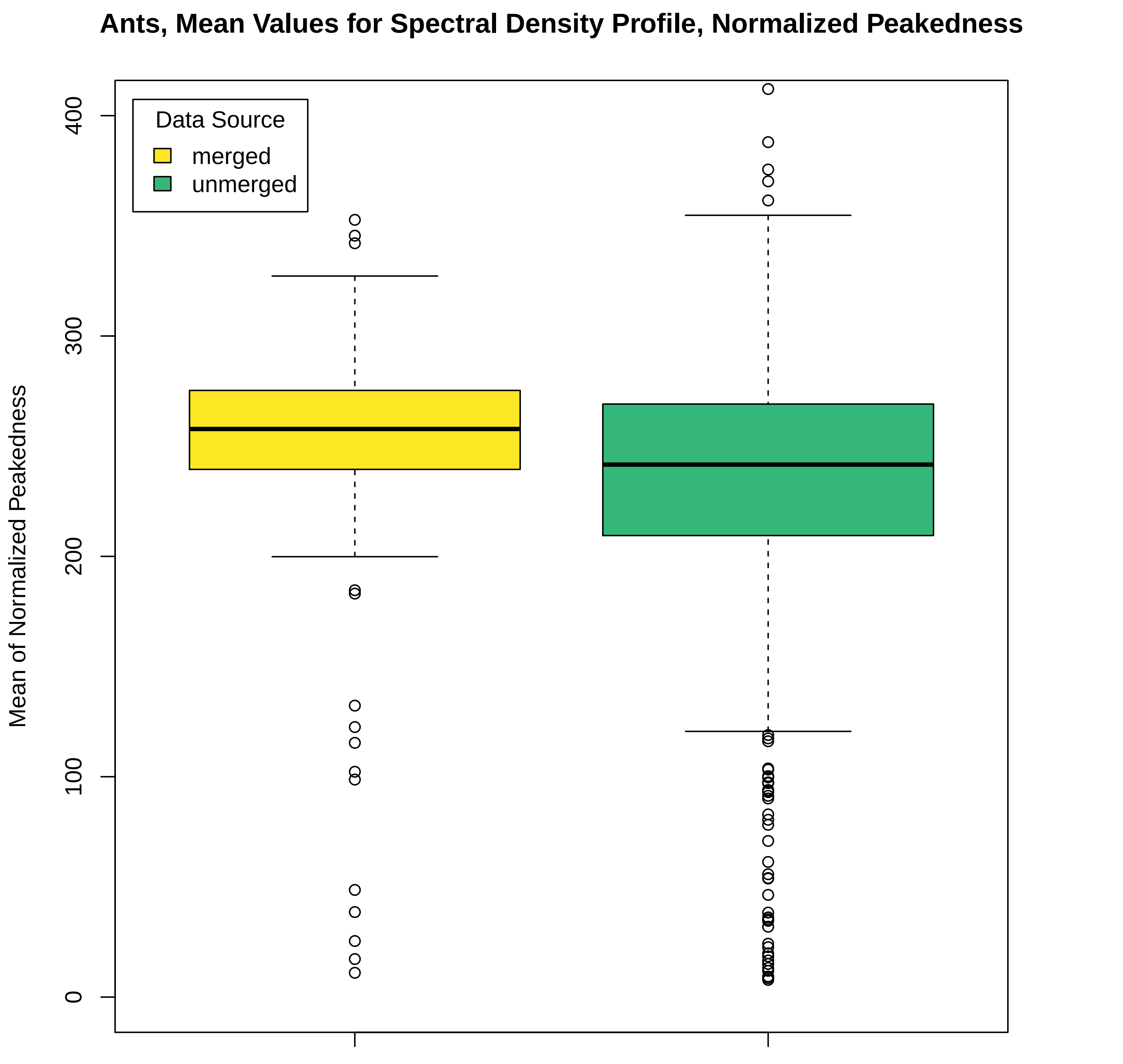

### Spectral_Density_Profile_Normalized_Peakedness_Birds.pdf

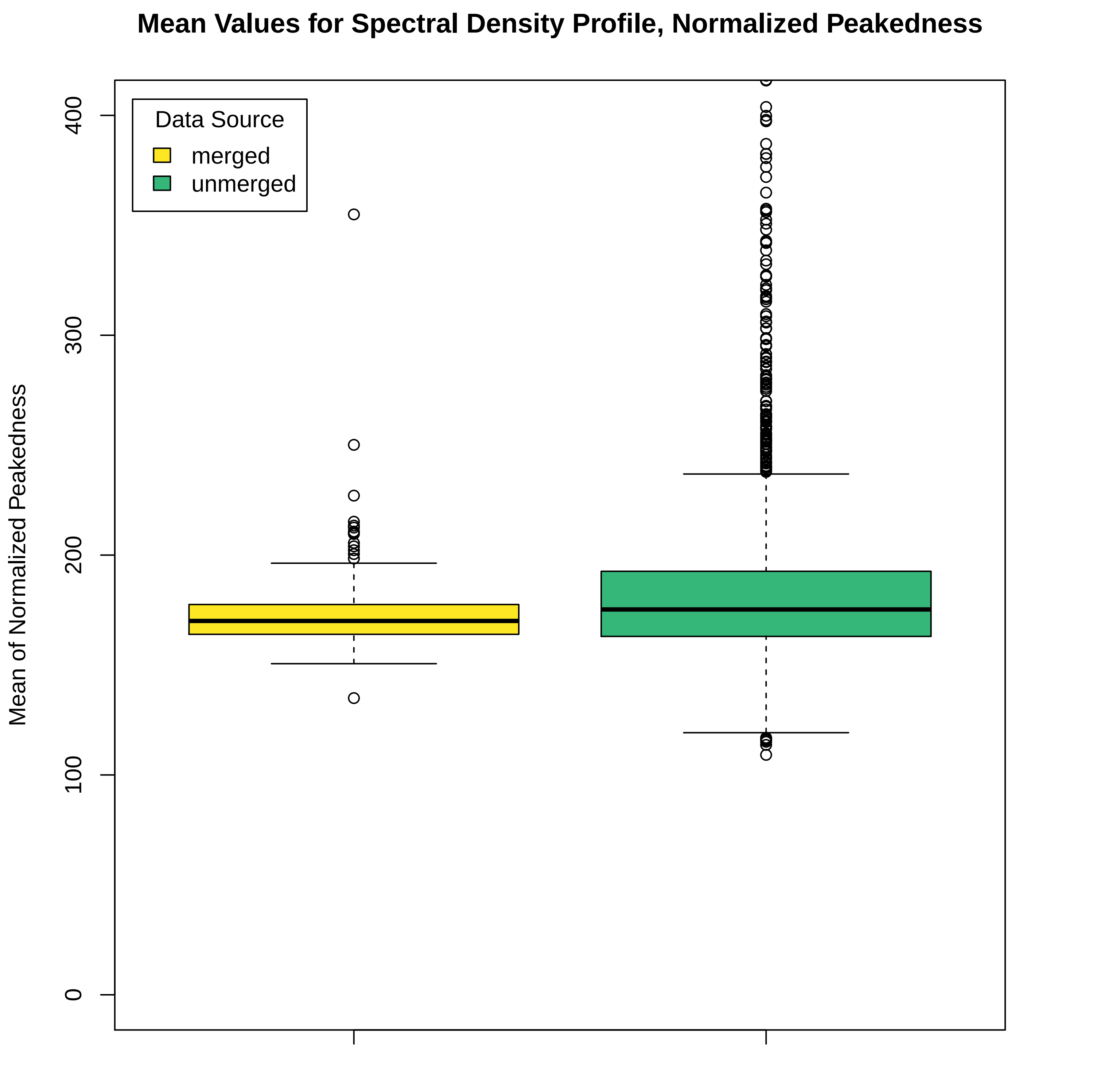

### Spectral_Density_Profile_Normalized_Peakedness_Fish.pdf

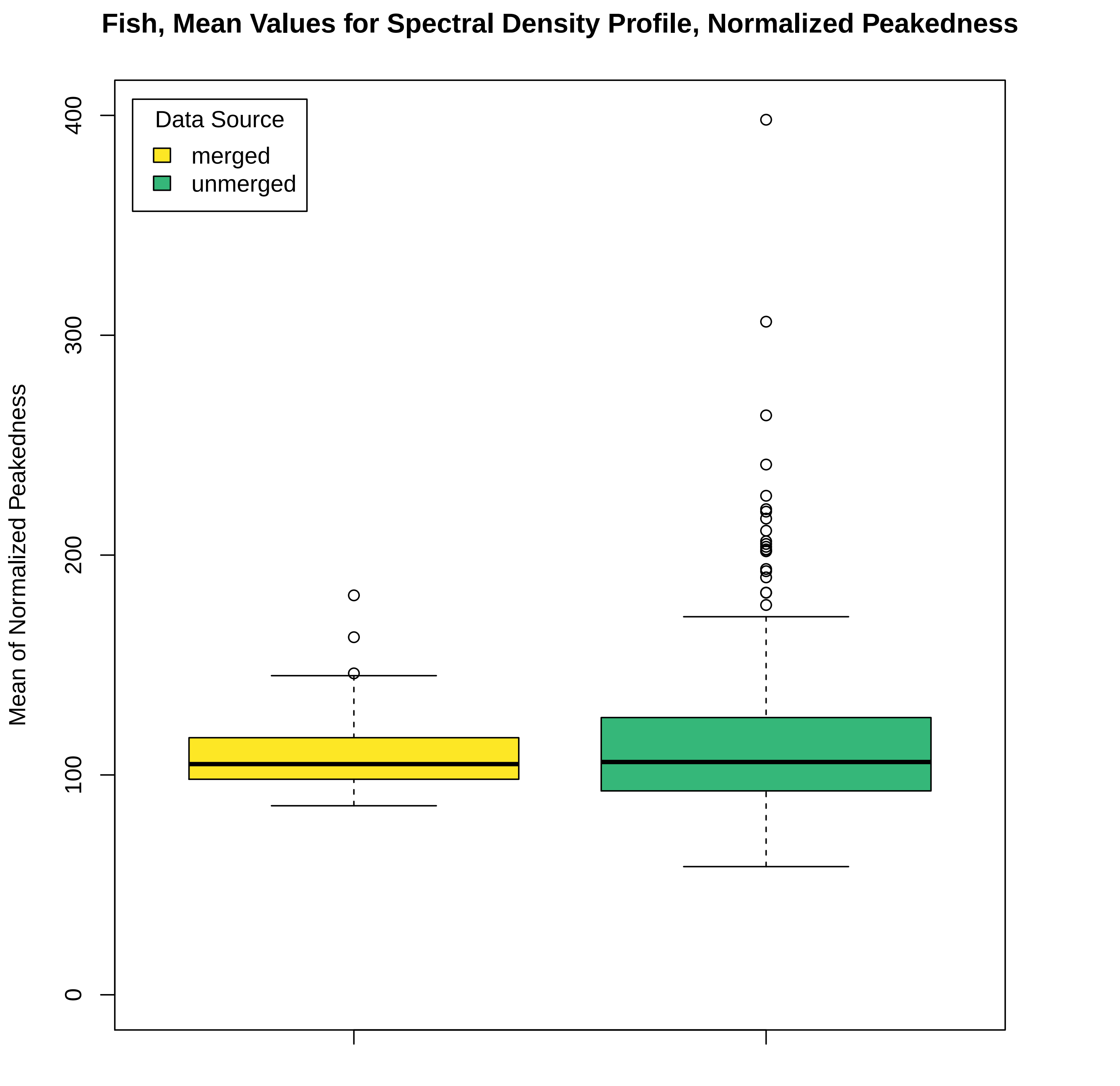

### Spectral_Density_Profile_Normalized_Peakedness_Mammals.pdf

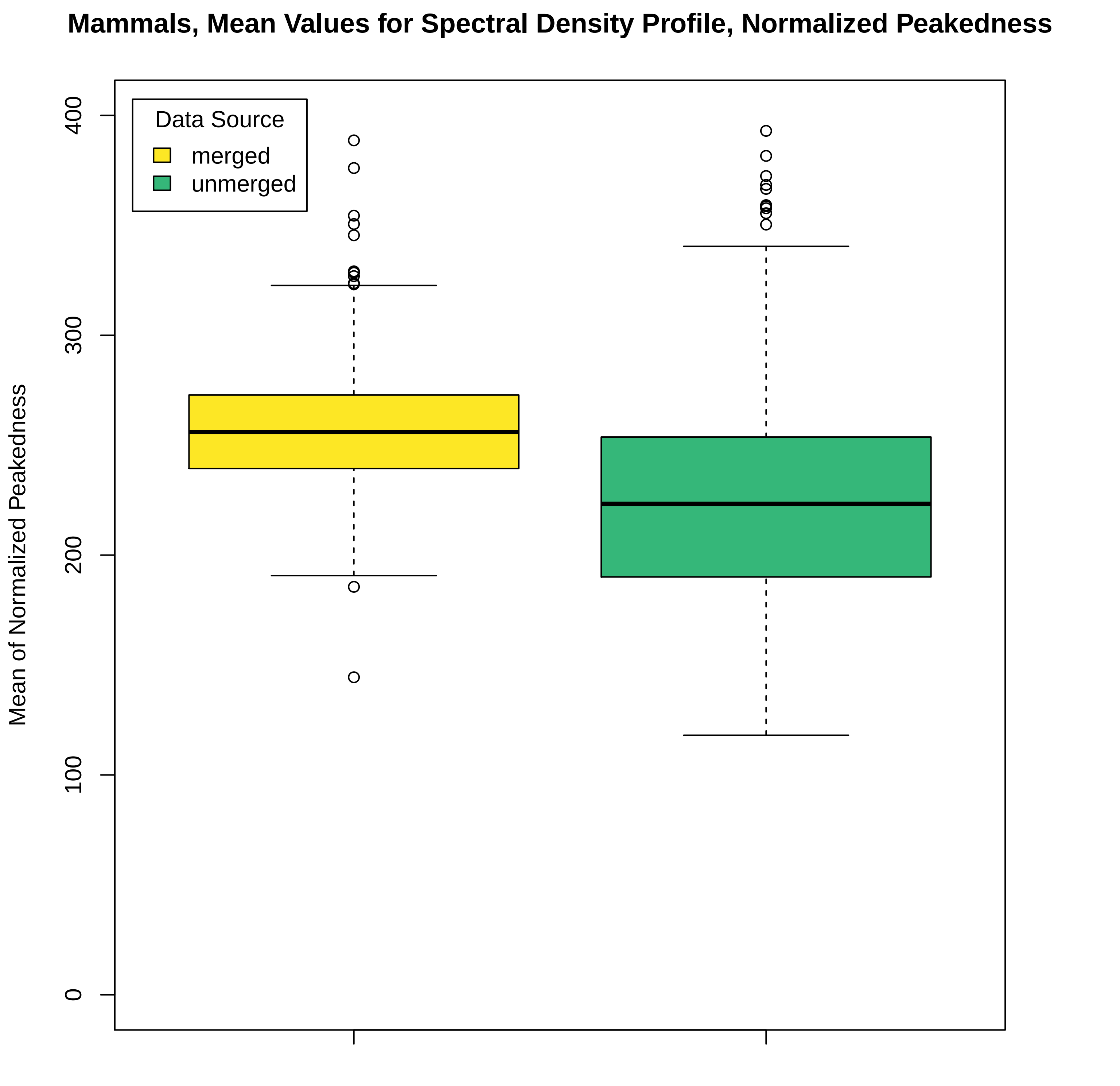

### Spectral_Density_Profile_Normalized_Peakedness_Weevils.pdf

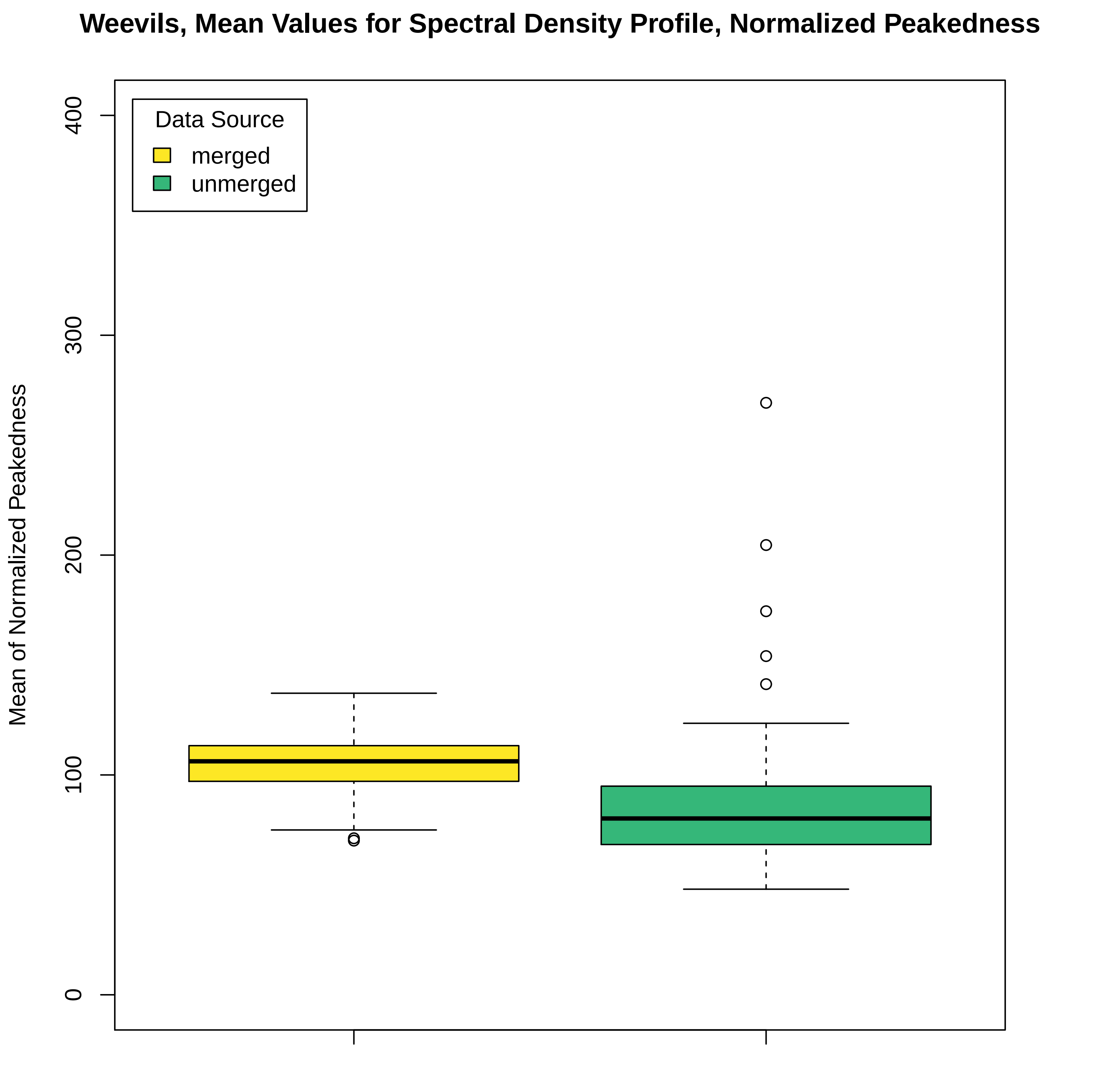
